# Loop extrusion dynamics and cooperative activation independently regulate enhancer-driven transcription

**DOI:** 10.64898/2026.05.06.723196

**Authors:** Elias T. Friman, Hannah J. Jüllig, Grace Alston, Luciana Gomez-Acuña, Shelagh Boyle, Milena Flankova, Alba Muñoz Grande, Giovanna Weykopf, Anita Mann, Iain Williamson, Laura A. Lettice, Wendy A. Bickmore

## Abstract

How enhancers transmit their activation signals to target promoters is a fundamental unanswered question. Gene activation by distal enhancers is affected by the three-dimensional organisation of chromatin driven by cohesin-mediated loop extrusion, but how the dynamic loop extrusion cycle shapes transcriptional outcomes is not fully understood. In addition, genes are typically regulated by multiple enhancers and cases of cooperativity between enhancers have been observed that depend on the relative position of enhancers and gene. The reason for such context-dependent cooperativity is unknown, and whether it is linked to loop extrusion has not been explored. Here, we use synthetic gene activation to show that NIPBL depletion, which decreases loop extrusion rate, reduces long-range activation in a dose-dependent manner. In contrast, WAPL depletion, which reduces cohesin turnover, can both increase and decrease distal gene activation depending on the presence of adjacent CTCF binding sites. These effects on gene activation correlate with the impact on chromatin contacts. Depletion of the co-activator BRD4 also affects long-range activation, but not by changing chromatin contacts. Synthetic activation from two locations results in super-additive responses, which can be predicted by an independently determined response model. These predictions hold for dual activation from proximal and/or distal sites, and with perturbed NIPBL, WAPL, or BRD4. Distal activation therefore appears to be equivalent in nature to proximal activation, and context-dependent cooperativity can arise simply from different level of activation inputs operating on a non-linear response function.

## Introduction

Hundreds of thousands of enhancers, each composed of clusters of transcription factor (TF) binding sites, control tissue-specific gene activation. Most genes are controlled by many enhancers, which may be active in different or overlapping contexts [1,2]. How the multitude of enhancers drive overall gene activity is poorly understood.

Combinations of enhancers can result in different types of non-additive effects. For example, two enhancers can be entirely redundant (deletion of one enhancer has no impact on transcription), or obligately cooperative (deletion of one enhancer abrogates transcription). A broad range of enhancer-enhancer combinatorial effects have been reported, and cooperativity can depend on the relative genomic position of the enhancers and gene [3–12]. What drives such context-dependent cooperativity is not known.

Many genes important in development are controlled by exceptionally long-range enhancers, several hundred kilobases (kb) or over one megabase (Mb) from their promoter, and even embedded in the introns of neighbouring genes [13,14]. This includes *Shh*, which encodes a developmentally critical morphogen and is regulated by multiple distal enhancers in various tissues, including the limb-specific ZRS enhancer, 850kb in mouse and 1Mb in human away from the gene (Fig. 1A) [15,16]. Another example is *Sox9*, encoding a developmental TF with roles in sex determination, skeletal and craniofacial development, which has several validated enhancers at up to 1.2Mb from the promoter in the mouse genome - 1.4Mb in human [17]. Such distal regulation must rely on communication between enhancer and promoter, but the underlying mechanisms remain largely unknown. Furthermore, whether there is a fundamental difference in how a gene responds to promoter-proximal and distal activation signals is an unanswered question [18–20].

**Figure 1.**
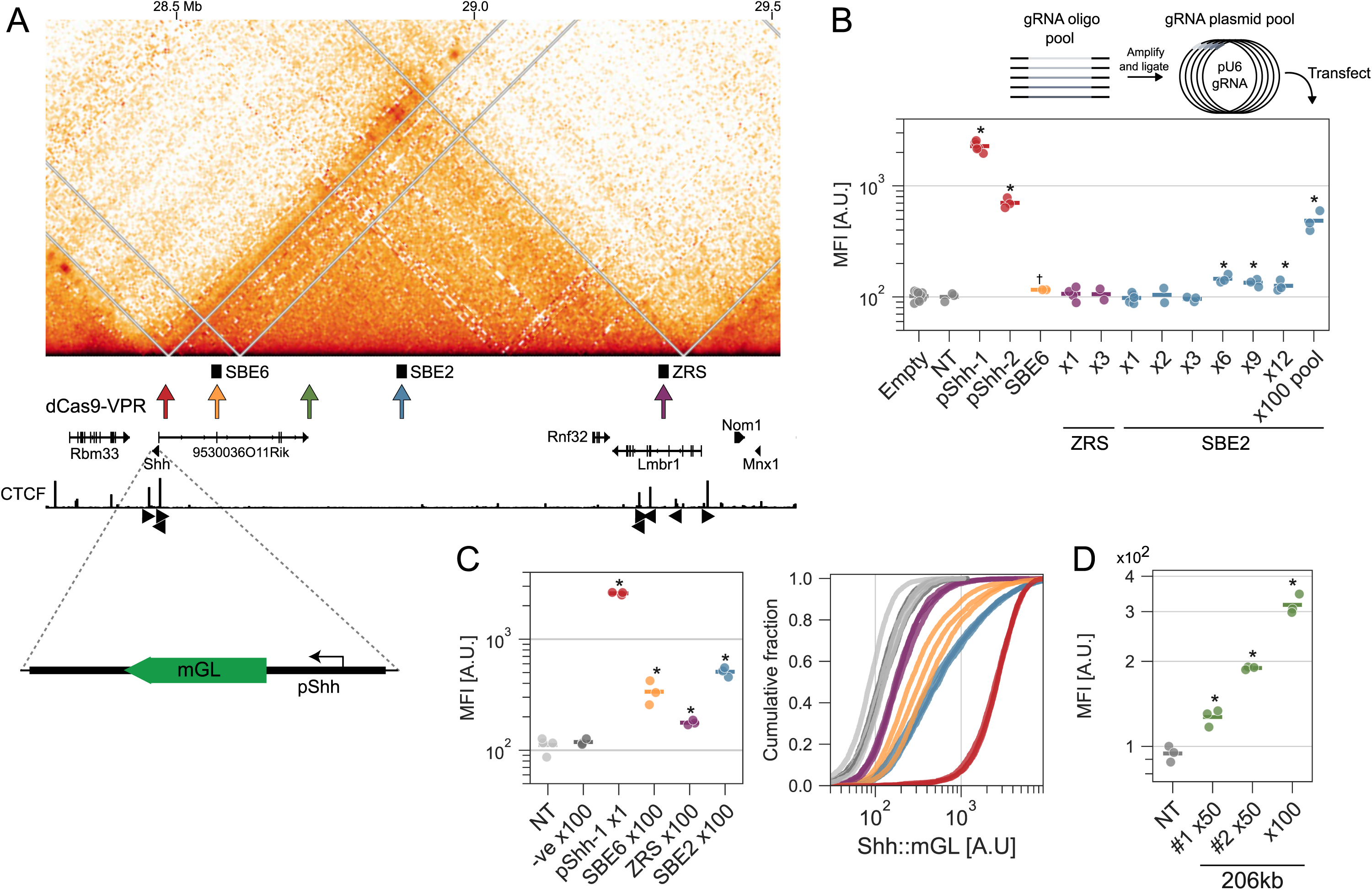
Distal activation of a *Shh* reporter. A) Micro-C data in WT mESCs [137] for the *Shh* TAD on chr5 (mm10 coordinates). dCas9-VPR arrows represent regions targeted by gRNAs, and black rectangles a subset of endogenous *Shh* enhancers in other tissues (SBE6, SBE2, ZRS). Below: CTCF ChIP-seq signal [138] and orientation of CTCF motifs (black arrowheads). The Shh::mGL reporter construct is illustrated at the bottom. B) Top: schematic for cloning gRNA pools. Below: Shh::mGL cells were transfected with dCas9-VPR and guides targeting different locations and transfected cells analysed for median mGreenLantern fluorescence (MFI) in arbitrary units [A.U.] using flow cytometry. Empty represents no guide, while NT represents a non-targeting guide. pShh-1/2 target the promoter, while SBE6/ZRS/SBE2 target regions in the *Shh* TAD (see A). Each dot represents one replicate. C) Left: As in B, but for cells transfected with dCas9-VPR and NT or gRNA pools targeting the different regions. -ve is a region 2 Mb downstream of *Shh* in another TAD. Right: Cumulative distribution of single-cell fluorescence values used to derive the median, with colour coding as in the left panel. Each line is a replicate. D) As in B: but for cells transfected with dCas9-VPR and NT or different gRNA pools targeting a site 206kb upstream of *Shh*. Values in B-D were statistically compared to Empty/NT with t-tests, BH-corrected, and adjusted p-values <0.1 (†) and <0.05 (*) shown (see Supplemental table 1 for statistics and replicate numbers).

Models of enhancer-promoter communication where signals are transmitted linearly along the chromatin fibre are unlikely to act at the most long-range enhancers, and there is limited evidence for such mechanisms [13,21]. In contrast, much evidence - including forcing enhancer-promoter contacts and correlations between contacts and enhancer influence, support a functional role for 3D genome organisation (contacts assayed by chromatin conformation capture methods such as Hi-C and Micro-C) [22–24]. This could involve direct enhancer-promoter physical contact or more general proximity allowing for diffusion [13,21,25–27]. There is not always a correlation between gene transcription and increased enhancer-promoter contacts (or decreased distances) [28–30]. Indeed, activation often co-occurs with increased enhancer-promoter spatial distances [31–33], which does not preclude an importance for proximity but complicates the causal and temporal relationships.

A major factor driving 3D genome organisation is cohesin, a complex consisting of core subunits SMC1, SMC3, and RAD21 (SCC1) [34,35]. Cohesin is regulated by co-factors which sustain its two major functions, sister chromatid cohesion and loop extrusion. Loop extrusion is an ATP-dependent process that occurs throughout interphase. The cohesin molecular motor embraces DNA in a pseudo-topological manner driving chromatin movement that can bring distal genomic regions (up to ∼2Mb) on the same chromosome together [35,36]. The major sites for regulation of loop extrusion are in RAD21 where multiple co-factors can bind. This includes NIPBL (SCC2) together with its partner protein MAU2 (SCC4), which are required for loop extrusion and may have a role in cohesin loading [37–41]. Depletion of RAD21 or NIPBL in mammalian cells leads to a dramatic reorganisation of the 3D genome. Contacts within self-interacting regions called topologically associated domains (TADs) are lost [36,42,43], and imaging shows increased distances between regions within, or at the border of, TADs [44–46].

TADs are formed by blocking of loop extrusion via CTCF, a zinc finger TF that binds to chromatin and can stall cohesin in a directional manner, dependent on the relative orientation of its DNA motif and oncoming extruding cohesin [47–49]. Each TAD border typically has at least one CTCF motif facing “inwards“ [36,47,48]. WAPL promotes cohesin unloading and cells depleted of WAPL have larger TADs and longer and stronger loops between convergently orientated CTCF sites [39,40,50,51]. The increased cohesin lifetime upon WAPL loss eventually leads to so-called “vermicelli” chromosomes where stacked chromatin-bound cohesin molecules can be seen by microscopy [39,52]. Cohesin regulators thus create a dynamic cycle of loading, extrusion, stalling, and unloading. In summary, perturbation of factors promoting (e.g. RAD21, NIPBL) or impeding (e.g. WAPL) loop extrusion have dramatic and apparently counteracting effects on genome organisation.

Genes and their enhancers tend to be localised in the same TAD [53,54]. This is particularly clear for developmental genes, including *Shh* and *Sox9*, where the gene is at the edge of a large TAD containing multiple enhancers, some very distal [16,17]. Such regulatory and structural coincidence led to hypotheses in which chromatin contacts promoted by loop extrusion within TADs, and suppressed between adjacent TADs, would positively or negatively influence enhancer-promoter communication [34,54]. Loop extrusion would thus be a major indirect regulator of transcription via its impact on enhancer-promoter contacts. However, perturbing cohesin leads to relatively minor transcriptional changes and enhancer-promoter contacts appear to be largely unaffected [36,42,55–60]. More specifically, synthetic activation by TF recruitment, genetic enhancer insertions, or global transcriptional analysis upon perturbation have shown that distal enhancers in mammalian cells are sensitive to cohesin [44,61–65]. How altering loop extrusion dynamics or cohesin turnover affects enhancer-driven transcription, and enhancer cooperativity, requires further investigation.

In this study we activate an otherwise silent endogenous promoter from different genomic locations in mouse embryonic stem cells (mESCs), achieved by targeted recruitment of synthetic transcription factors to distal regions without genetically modifying the locus. We combine this with rapid depletion of NIPBL and WAPL to test the role of loop extrusion dynamics across different enhancer-promoter distances. By activating from multiple genomic locations simultaneously, we also test the role of loop extrusion in driving cooperativity. Our results suggest that the dynamic loop extrusion cycle affects distal enhancers, and that loop extrusion and cooperativity are independent and important aspects of activation from enhancers.

## Results

### Long distance gene activation requires large numbers of activators

Many genes are regulated by multiple enhancers, but their individual and combined influence, in any particular cell type, is often not known and difficult to test. Global measurements of transcription following perturbations are difficult to interpret without this knowledge. To circumvent these limitations, and to study distal gene activation and cooperativity, we aimed to activate an otherwise silent gene using targeted recruitment of activators to multiple locations, reading out gene activation using a fluorescent reporter. *Shh* is transcriptionally silent, repressed by Polycomb, with no evidence that any of its endogenous enhancers are active in mESCs (Suppl. Fig. 1A) [31]. Previously, we have used synthetic Transcription Activator Like Effectors fused to multimers of VP16 (TALE-VP64) as programmable TFs targeted to known *Shh* enhancers in mESCs, assaying induced transcription using nascent RNA-fluorescence in situ hybridisation (RNA-FISH) [44]. However, TALEs are not amenable to multiplexing activators at many diverse sites, and RNA-FISH is a laborious assay.

To quantify transcription at the *Shh* promoter at higher throughput, we homozygously replaced the *Shh* gene (deleting introns and exons but retaining the promoter, and the 5’ and 3’ untranslated regions of the mRNA) in mESCs with mGreenLantern (Fig. 1A, Suppl. Fig. 1B-D), which encodes for a bright green fluorescent protein detectable using flow cytometry [66]. Catalytically dead Cas9 (dCas9)-activator fusions (CRISPRa [67]) have been used to drive gene expression using single guide RNAs (gRNAs), and a composite of activation domains VP64, p65 and Rta (VPR), has been shown to be one of the strongest synthetic activators in multiple contexts [68,69]. However, activation efficiency falls off rapidly with distance of gRNA binding sites from the promoter, with strongly diminished effect even at tens of kb [68,70–72]. Longer-range activation (>100kb) has been demonstrated, but this was in tissues or cell types where the enhancer is already active or with multiple gRNAs [70,73,74].

We were able to induce high levels of the *Shh* mGreenLantern (Shh::mGL) reporter following transient transfection of dCas9-VPR and a gRNA targeted to the *Shh* promoter (pShh), which peaked two days after transfection (Fig. 1B, Suppl. Fig. 2A). There was no background reporter activity without transfection and negligible activity for dCas9-VPR targeted with no guide (empty) or non-targeting (NT) guides (Suppl. Fig. 2B). However, we could not detect activation using single guides targeting the SBE6 (96kb), ZRS (848kb), or SBE2 (409kb) enhancers upstream of *Shh* (Fig. 1B), nor to regions closer to *Shh* (Suppl. Fig. 2C). We also could not boost activation by combining a gRNA to the promoter and one to the distal sites, as has been suggested (Suppl. Fig. 2D) [71]. This may be because the activation signal from a single distally bound dCas9-VPR is too weak to activate a gene across larger distances, or because of dCas9-VPR failing to bind at target sites due to, for example, nucleosome occlusion [75]. Increased distal activation efficiency has been achieved by targeting dCas9-VPR with multiple (up to six) gRNAs from sites up to ∼150kb away from a promoter [70]. Indeed, we could detect some Shh::mGL activation by targeting dCas9 to SBE2 with six gRNAs (Fig. 1B). However, the level of activation was very low compared with that achieved by targeting pShh and below what would be experimentally tractable for quantitative comparisons. This was not increased using 9 or 12 gRNAs. Therefore, to test if we could brute-force activation, we devised a cloning strategy to generate pools of plasmids with 100 unique gRNAs (one per plasmid) to target dCas9-VPR to a ∼1kb region at SBE2. This resulted in substantial activation of Shh::mGL (Fig. 1B).

We then designed x100 gRNA pools targeting SBE6 and ZRS, which also drove reporter activation but to lower levels than SBE2, despite SBE6 being closer to *Shh* in the genome (Fig. 1C, Suppl. Fig. 2E), suggesting variability in gRNA pool efficiencies. A negative control x100 gRNA pool targeting a site 2Mb away from *Shh* in another TAD did not activate the reporter (Fig. 1C).

To test pool-to-pool variability in activation efficiency independently of distance, we designed two adjacent pools of x50 gRNAs targeting a site 206kb upstream of *Shh* (Fig. 1D, Suppl. Fig. 2F). These activated to different extents, and both less than the combined x100 gRNAs. Activation between different guide pools can therefore not be directly compared because of varying, and unknown, efficiencies. Our ability to activate pShh from an arbitrary and apparently inert genomic locus at 206kb, and from endogenous enhancers which are not active in mESCs up to 848kb, suggests a pre-existing active or primed enhancer is not required for synthetic activation, in agreement with previous studies [70,72].

gRNAs are not equally represented in the pools (Suppl. Fig. 2G) and transfection is estimated to lead to thousands, or tens of thousands, of plasmids per cell [76,77]. Activation is therefore unlikely achieved by a few “jackpot” guides in the pool but rather the combined activity of multiple guides. We therefore chose to use x100 gRNAs for all future experiments.

Our results show that, by recruitment of multiple molecules of dCas9-VPR to a locus using large numbers of gRNAs, we can activate gene expression at a Polycomb-repressed gene (*Shh*) across very large genomic distances (up to 848kb). To our knowledge this is the longest-range gene activation that has been achieved by dCas9-VPR.

### NIPBL depletion reduces activation from distal sites in a dose-dependent manner

We, and others, have previously shown that cohesin depletion by inducible degradation of RAD21 particularly affects on gene activation from long-range enhancers, with more proximal enhancers apparently unaffected [44,61,63]. Though originally considered as only required for loading of cohesin onto chromatin, recent studies suggest that reduced NIPBL levels have a more profound effect on extrusion rate rather than the amount of cohesin-loaded [39–41]. Therefore, it is unclear whether there would be the same relationship between enhancer-promoter genomic distance and NIPBL depletion as was seen for RAD21 at the *Shh* locus.

To test this, we generated homozygous endogenous N-terminal fusions of FKBP12^F36V^-HaloTag to NIPBL (henceforth dTAG-NIPBL) in the Shh::mGL reporter line (Supplemental Fig. 3A-B). This enabled rapid (within 1 hour) depletion of NIPBL, to levels of a few percent of residual protein, following addition of dTAGV-1 (Supplemental Fig. 3C-D). Unlike RAD21 depletion, which leads to mitotic failure after a few hours in mESCs [78], likely due to problems with sister chromatid cohesion, depletion of NIPBL by 48 hours of dTAGV-1 treatment resulted in only a minor but statistically significant effect on the cell cycle (Supplemental Fig. 3E). As shown previously, long-term (7 day) NIPBL depletion led to loss of mESC pluripotency (Supplemental Fig. 3F) [79].

We used Region Capture Micro-C (RCMC) [80] and DNA-FISH to assess the effect of NIPBL depletion on chromatin organisation across the *Shh* TAD. After 24 hours of dTAGV-1 treatment there was a decrease in contacts within the TAD (Fig. 2A). Comparison to published Micro-C data shows that short-term RAD21 depletion results in a slightly greater loss of contacts at distances <1Mb than NIPBL depletion (Fig. 2B). This suggests that NIPBL depletion in our cell line leads to strongly reduced, but not a complete loss of, loop extrusion. DNA-FISH showed increased spatial distances between probes across the TAD, comparable to that seen with short-term RAD21 depletion (Fig. 2C, Suppl. Fig. 3G).

**Figure 2.**
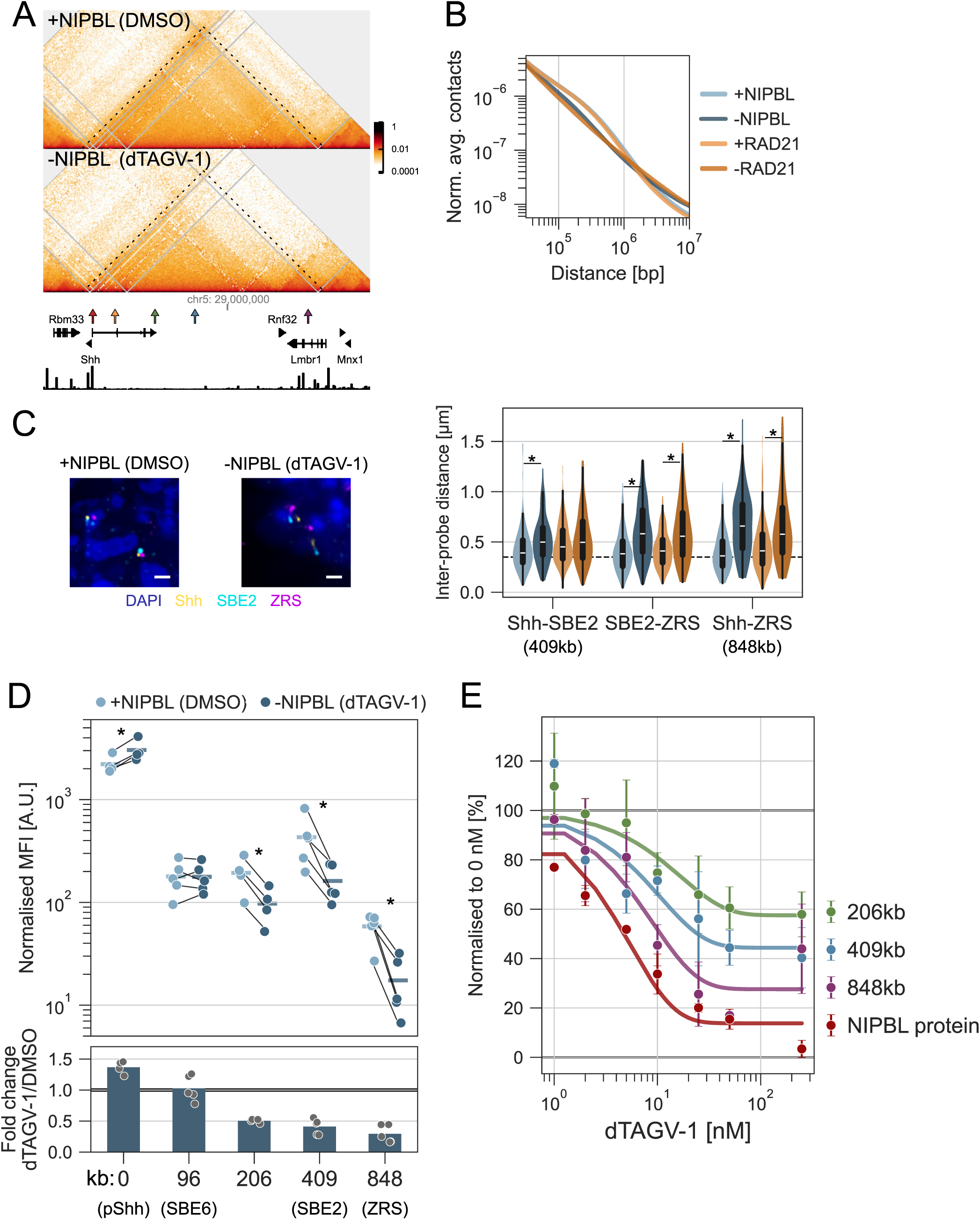
NIPBL depletion decreases distal activation in a dose-dependent manner. A) Region Capture Micro-C (RCMC) data for dTAG-NIPBL cells treated with DMSO (top) or 250 nM dTAGV-1 (bottom) for 24 hours. Colour bars show the scaling of normalised contacts. The dashed lines outline the *Shh* TAD. Coordinates are in mm10. Shown underneath are genes and CTCF ChIP-seq signal [138]. B) P(s) curve derived from RCMC data for dTAG-NIPBL cells treated with DMSO or dTAGV-1, and Micro-C data from RAD21-AID cells [57] untreated or treated with Auxin. Average contacts at a given genomic separation (bp) were normalised to the area under the curve. C) Left: Representative DNA-FISH hybridisation signals for Shh, SBE2 and ZRS in dTAG-NIPBL cells treated with DMSO (left) or dTAGV-1 (right) for 24 hours. Scale bar = 1 micron. Right: violin plots show the measured inter-probe distances (µm) between *Shh*-SBE2, SBE2-ZRS, *Shh*-ZRS dTAG-NIPBL cells depleted for NIPBL (DMSO vs dTAGV-1) for 24 hours or RAD21-AID cells depleted of RAD21 (untreated versus auxin) for 6 hours. Boxes show interquartile range (Q1-Q3) and whiskers show remaining data range. Dotted line indicates 350 nm. D) Top: Mean mGreenLantern fluorescence (MFI), normalised by subtracting the value of NT-transfected cells, in dTAG-NIPBL cells transfected with dCas9-VPR and different gRNA or gRNA pools at different distances from the *Shh* promoter (x axis) and treated with DMSO or 250 nM dTAGV-1 for 48 hours. Bottom: fold-change between dTAGV-1 and DMSO, with points and bars showing individual replicates and the mean, respectively. E) dTAG-NIPBL cells treated with different concentrations of dTAGV-1 for 48 hours and either stained for HaloTag-JF646 (NIPBL protein) and fluorescence measured using flow cytometry, or transfected with dCas9-VPR and different gRNA pools and mGreenLantern fluorescence measured using flow cytometry. Values were min-max normalised and a decay function fitted to the mean values. Error bars represent the standard deviation. Values in C were statistically compared between dTAGV-1 and DMSO with Mann-Whitney U tests and BH-corrected. Values in D were statistically compared between dTAGV-1 and DMSO with paired t-tests and BH-corrected. Adjusted p-values <0.1 (†) and <0.05 (*) are shown (see Supplemental table 1 for statistics and replicate numbers).

To determine how NIPBL depletion affects the ability of enhancers to activate the *Shh* reporter, we transfected dTAG-NIPBL cells with dCas9-VPR and a gRNA targeting pShh (0kb) or x100 gRNA pools targeting different distal locations in the *Shh* TAD. Simultaneously, we treated cells with DMSO or dTAGV-1, and measured reporter fluorescence after 48 hours by flow cytometry, normalising by subtracting the signal from the NT guide. NIPBL depletion had no effect on activation by dCas9-VPR targeted by a pool of gRNAs 96kb upstream of the Shh promoter, but there was a distance-dependent effect beyond that (Fig. 2D, Suppl. Fig. 3H). Reporter activation decreased upon activation from large distances (206-848kb) after NIPBL depletion, consistent with our previous result using a RAD21 degron [44]. NIPBL depletion slightly increased activation from a promoter targeted dCas9-VPR, as has been seen in other studies [65], but the mechanistic basis for this is unclear.

We wondered if some of the residual activation from the most distal location (ZRS, 848kb), upon NIPBL depletion could be due to a loop extrusion-independent mechanisms of enhancer-promoter spatial communication, as has been suggested in the limb where ZRS is active [81]. To this end, we deleted ZRS and activated from a ZRS-adjacent upstream region 4kb closer to *Shh* (Suppl. Fig. 4A). ZRS deletion had a minor but significant effect on the level of activation in DMSO-treated control cells but not in the dTAGV-1 treated cells depleted of NIPBL, suggesting that in mESCs the ZRS itself has a minor contribution to chromatin contacts with *Shh* (Suppl. Fig. 4B).

NIPBL haploinsufficiency is the most common cause of the developmental disorder Cornelia de Lange syndrome (CdLS). Despite heterozygous loss of function mutation in *NIPBL* in most cases of severe CdLS [82], levels of *NIPBL* mRNA are reported to not drop below 60% in CdLS cells compared with controls, suggesting some degree of transcriptional compensation [83]. To assess how long-range activation is affected by NIPBL dose, we assayed the effects on reporter activation in the presence of partially depleted NIPBL levels, achieved by dTAGV-1 titration (Fig. 2E). Levels of reporter activation from distances of 206-848kb scaled with the level of NIPBL, with depletion of ∼30% NIPBL leading to detectably lower activation. A NIPBL-dose response on chromatin structure was also detected by RCMC (Suppl. Fig. 4C). These results show that levels of NIPBL affect the level of loop extrusion, which in turn affects distal enhancer influence.

### WAPL depletion can both increase and decrease distal activation

Cohesin typically resides on chromatin for a few tens of minutes [84,85], estimated to be enough time to extrude several hundreds of kb of chromatin (at an estimated speed in the order of ∼0.5kb per second [36,39]). However, the residence time of cohesin on chromatin is dramatically increased, to hours, following depletion of WAPL, and larger extruded loops therefore become more frequent [50,51,86].

To test the effect of reduced cohesin unloading and extended loop extrusion on distal gene activation, we generated endogenous homozygous C-terminal FKBP12^F36V^-HaloTag fusions to WAPL (henceforth WAPL-dTAG) in the Shh::mGL reporter background (Suppl. Fig. 5A-D). WAPL depletion did not perturb the cell cycle within 48 hours but led to differentiation of mESCs upon long-term (7 day) dTAGV-1 treatment, as seen previously (Suppl. Fig. 5E-F) [86].

RCMC showed that 3D chromatin conformation within the *Shh* TAD was more subtly affected by depletion of WAPL than by NIPBL, but with a slight increase in contacts between *Shh* and ZRS (Fig. 3A, arrowhead). WAPL depletion globally slightly decreased contacts below and increased contacts above ∼500kb (Fig. 3B). Despite reduced intra-TAD contacts as assayed by RCMC, DNA-FISH showed no significant change in intra-TAD distances, but did show a statistically significant decrease in the average *Shh-*ZRS (848kb) distances in one of two replicates (Fig. 3C, Suppl. Fig. 5G). These results are in line with reduced loop-extruding complexes within the TAD, and some degree of increased cohesin stalling at CTCF sites at the *Shh* TAD boundaries near *Shh* and ZRS.

**Figure 3.**
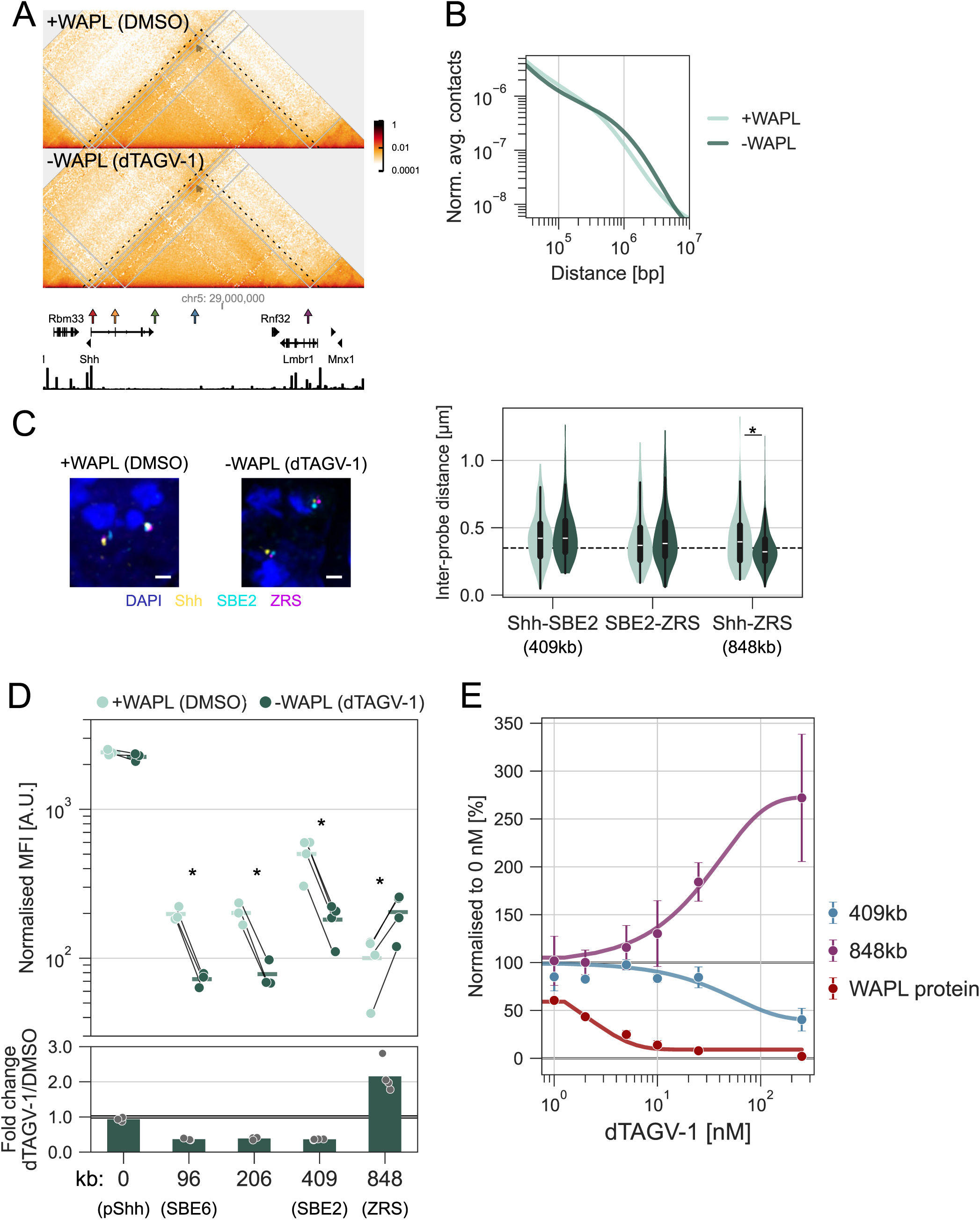
WAPL depletion increases or decreases distal activation in a position-dependent manner. A) Region Capture Micro-C (RCMC) data for WAPL-dTAG cells treated with DMSO (top) or 250 nM dTAGV-1 (bottom) for 24 hours. Colour bars show the scaling of normalised contacts. The dashed lines outline the *Shh* TAD. Arrows show contacts between the regions harbouring *Shh* and ZRS. Coordinates are in mm10. Shown underneath are genes and CTCF ChIP-seq signal [138]. B) P(s) curve derived from RCMC data for WAPL-dTAG cells treated with DMSO or dTAGV-1. Average contacts at the given genomic separation (bp) were normalised to the area under the curve. C) Left: Representative DNA-FISH hybridisation signals for Shh, SBE2 and ZRS in WAPL-dTAG cells treated with DMSO (left) or dTAGV-1 (right) for 24 hours. Scale bar = 1 micron. Right: violin plots show the measured inter-probe distances (µm) between Shh-SBE2, SBE2-ZRS, Shh-ZRS in WAPL-dTAG cells depleted for WAPL (DMSO vs dTAGV-1) for 24 hours. Boxes show interquartile range (Q1-Q3) and whiskers show remaining data range. Dotted line indicates 350 nm. D) Top: Mean mGreenLantern fluorescence (MFI), normalised by subtracting the value of NT-transfected cells, in WAPL-dTAG cells transfected with dCas9-VPR and different gRNA or gRNA pools at different distances from the Shh promoter (x axis) and treated with DMSO or 250 nM dTAGV-1 for 48 hours. Bottom: fold-change between dTAGV-1 and DMSO, with points and bars showing individual replicates and the mean, respectively. E) WAPL-dTAG cells treated with different concentrations of dTAGV-1 for 48 hours and either stained for HaloTag-JF646 (WAPL protein) and fluorescence measured using flow cytometry, or transfected with dCas9-VPR and different gRNA pools and mGreenLantern fluorescence measured using flow cytometry. Values were min-max normalised and a decay function fitted to the mean values. Error bars represent the standard deviation. Values in C were statistically compared between dTAGV-1 and DMSO with Mann-Whitney U tests and BH-corrected. Values in D were statistically compared between dTAGV-1 and DMSO with paired t-tests and BH-corrected. Adjusted p-values <0.1 (†) and <0.05 (*) are shown (see Supplemental table 1 for statistics and replicate numbers).

Unlike for NIPBL depletion, WAPL depletion had no effect on activation of pShh by a promoter-targeted dCas9-VPR but decreased the ability to activate with the 96, 206, and 409kb-targeting oligo pools (Fig. 3D, Suppl. Fig. 5H). Strikingly, WAPL depletion resulted in an increased ability to activate the reporter from the most distal location (848kb, at ZRS), which is near CTCF sites. This is consistent with the increased contacts between the *Shh* TAD boundaries and an accumulation of stalled cohesin there in the absence of the cohesin unloader. Compared to NIPBL, titrating the level of WAPL showed a less dose-dependent relationship to reporter activation, with activation affected mainly after WAPL levels approached <20% (Fig. 3E). These data show that reducing cohesin turnover can both decrease and increase enhancer influence, depending on whether the activation occurs proximal to CTCF sites (or TAD boundaries) or not.

### Cohesin perturbation affects distal Sox9 activation

To test the effects of NIPBL and WAPL depletion on another distally regulated gene, we generated polyclonal reporters for *Sox9* transcription (Sox9-P2A-mScarlet3) in the dTAG-NIPBL and WAPL-dTAG cell lines (Fig. 4A, Suppl. Fig. 6A). Although a Polycomb target (Suppl. Fig. 6B), we found that, unlike for *Shh,* the *Sox9* reporter was active above background in mESCs. To mitigate precocious differentiation, we added GSK-3 and MEK inhibitors (2i) to the culture medium. We used dCas9-VPR to activate *Sox9* from its promoter with a single gRNA or from three distal locations with x100 gRNA pools. These include a location ∼1.2Mb upstream at the other end of the TAD, where an endogenous enhancer cluster active in neural crest resides [17], and two arbitrary locations at 715kb and 98kb upstream. Both the 1.2Mb and 715kb locations are near several CTCF peaks, while the 98kb location is not (Suppl. Fig. 6B).

**Figure 4.**
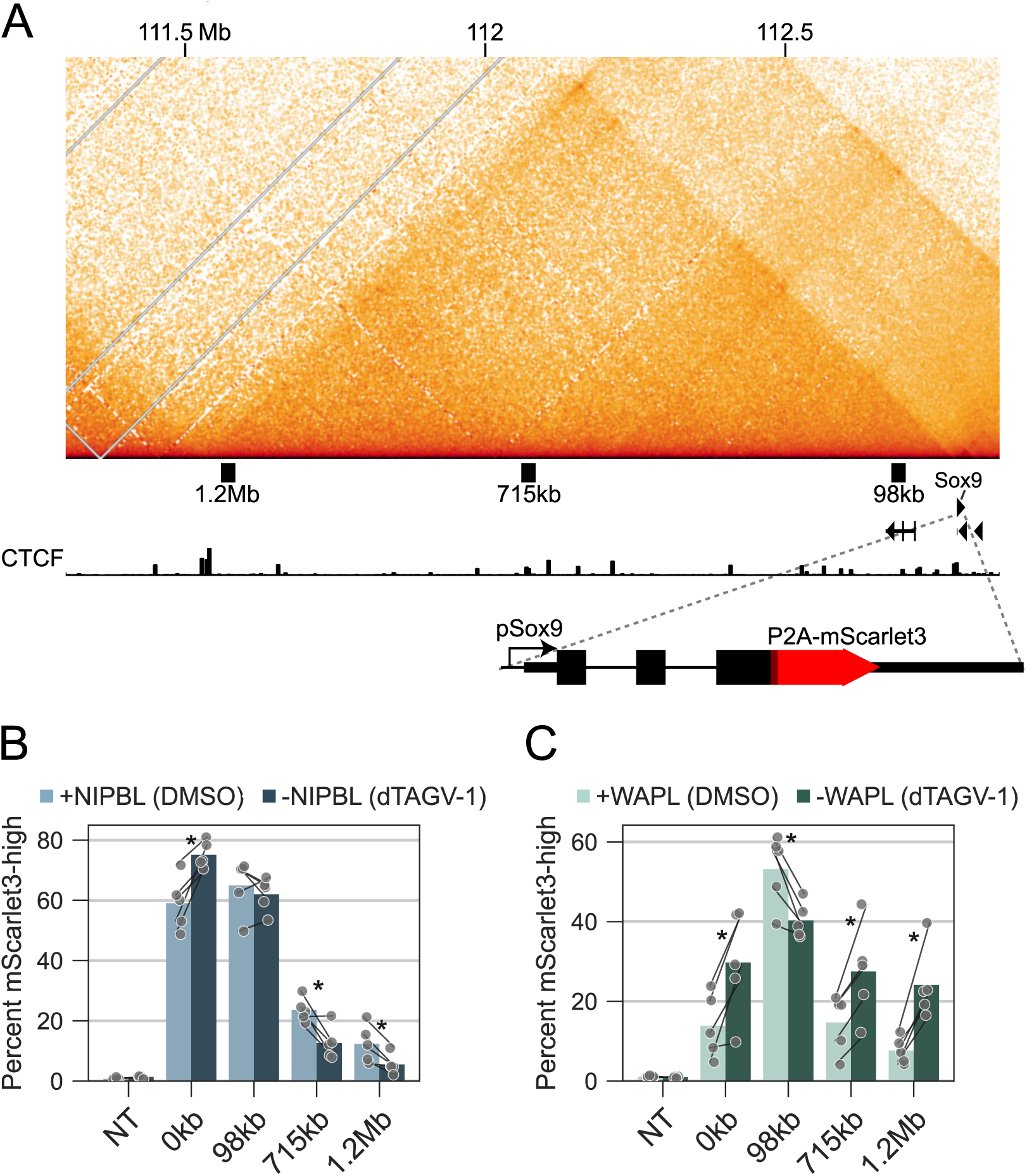
*Sox9* distal activation is affected by NIPBL and WAPL depletion. A) Micro-C data in WT mESCs [137] at the *Sox9* TAD on chr11 (mm10 coordinates). Shown underneath are genes and CTCF ChIP-seq signal [138]. Black rectangles represent regions targeted by dCas9-VPR in subsequent experiments. The Sox9-P2A-mScarlet3 reporter construct is illustrated at the bottom. B) Percentage of mScarlet3-high cells, defined as the signal above the highest 1% in NT-transfected cells, for dTAG-NIPBL *Sox9* reporter cells transfected with dCas9-VPR and different gRNA or gRNA pools (x axis) and treated with DMSO or dTAGV-1 for 48 hours. C) As in (B) but for WAPL-dTAG *Sox9* reporter cells Values in B-C were statistically compared between dTAGV-1 and DMSO with paired t-tests and BH-corrected. Adjusted p-values <0.1 (†) and <0.05 (*) are shown (see Supplemental table 1 for statistics and replicate numbers).

Large variability in background Sox9-P2A-mScarlet3 activation in the NT condition, and a large spread of fluorescence values (Suppl. Fig. 6C), led to the median NT value sometimes being higher than the activation condition (Suppl. Fig. 6D-E). We attribute this to the polyclonal nature of the cells and background activity of the reporter, which may vary with cell state. To ensure we were measuring only synthetic activation, we dynamically thresholded the mScarlet3 signal to count the percent of cells with a level of fluorescent reporter above that of the top 1% of NT-transfected cells. Similar to the data at *Shh*, NIPBL depletion increased activation from the promoter, did not significantly affect activation at 98kb, but reduced reporter activation at 715kb and 1.2Mb (Fig. 4B). In contrast, WAPL depletion decreased activation at 98kb, and increased the activation at 715kb and 1.2Mb, consistent with the proximity to CTCF sites (Fig. 4C). We also found that WAPL depletion increased activation from the promoter, which was not expected. We speculate that this may be due to an indirect effect specific to *Sox9* activation, which leads to overexpression of a strong TF (Sox9) involved in differentiation. Nevertheless, these results are in broad agreement with our findings at the *Shh* locus, where NIPBL depletion reduces distal (>100kb) activation and the effect of WAPL depletion depends on the relative position of the targeted activators and CTCF sites.

### BRD4 affects distal activation independent of loop extrusion

*NIPBL* mutation is the most frequent cause of severe CdLS, with less frequent mutations in other cohesin components (*SMC1A, SMC3, RAD21*) or regulators (*HDAC8*) associated with less severe atypical CdLS [87]. Mutations in *BRD4* have also been associated with atypical CdLS [88,89]. BRD4 is a BET domain protein whose bromodomains (BDs) recognise acetylated histones and has been implicated in enhancer function [90,91]. Protein interactions between BRD4 and NIPBL have been detected [88,92] and reported non-additive effects of BD inhibition and RAD21 depletion on enhancer-driven gene activation have been suggested to indicate that BRD4 and cohesin operate in a shared pathway [93].

To test the impact of BRD4 reduction on distal gene activation, with or without cohesin perturbation, we partially depleted BRD4 using a low dose of the BET PROTAC MZ1, shown to specifically target BRD4 (Suppl. Fig. 7A) [94]. MZ1 had no impact on Shh::mGL reporter activation from a promoter-targeted dCas9-VPR, but it attenuated activation from two out of three distal activator pools in the *Shh* TAD, namely at 96kb and 409kb distance (Fig. 5A). Co-depletion of NIPBL (dTAGV-1) with BRD4 reduction was synergistic, with MZ1 treatment sensitising activation from 96kb and 409kb to NIPBL depletion. This is inconsistent with BRD4 operating through direct effects on loop extrusion. BRD4 degradation has been reported to reduce the association of NIPBL with chromatin and to weaken some TADs [92]. However, reanalysis of Hi-C data from BRD4-dTAG cells revealed no clear reduction in contacts in the *Shh* TAD upon BRD4 depletion alone, suggesting the reduced distal activation is not mediated by loss of chromatin contacts (Suppl. Fig. 7B) [92].

**Figure 5.**
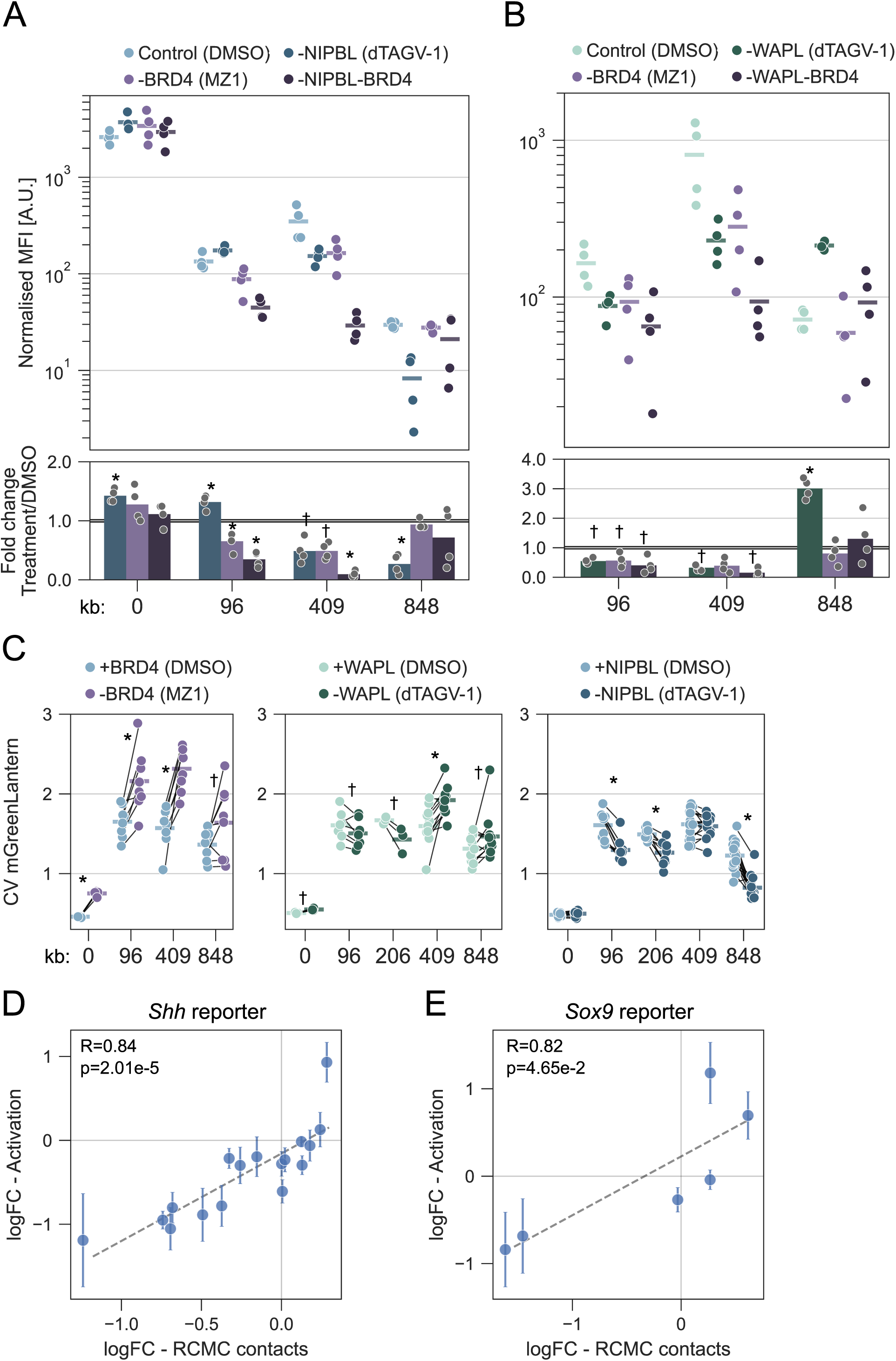
Effect of BRD4 depletion and cohesin perturbations. A) Top: mGreenLantern fluorescence, normalised by subtracting the value of NT-transfected cells, for dTAG-NIPBL cells transfected with dCas9-VPR and different gRNA or gRNA pools (x axis) and treated with DMSO, dTAGV-1, MZ1, or dTAGV-1 and MZ1 for 48 hours. Bottom: fold-change between treatment and DMSO. B) As in (A) but for WAPL-dTAG cells. C) Coefficient of variation (CV) of the mGreenLantern signal of the transfected population upon depletion of BRD4, WAPL, and NIPBL. D) Log fold-change in RCMC contacts between the *Shh* reporter and the activation site (96kb, 206kb, 409kb, 848kb) upon treatment (x axis), correlated against the log fold-change in *Shh* reporter activation upon treatment (y axis; conditions are dTAG-NIPBL DMSO/dTAGV-1 doses, and WAPL-dTAG DMSO/dTAGV-1). Error bars represent the standard deviation of activation data. E) Log fold-change in Micro-C contacts [39,57] between *Sox9* and the activation site (98kb, 715kb, 1.2Mb) upon treatment (x axis) correlated against the log fold-change in *Sox9* reporter activation upon treatment (y axis; conditions are dTAG-NIPBL DMSO/dTAGV-1 and WAPL-dTAG DMSO/dTAGV-1). Error bars represent the standard deviation of activation data. Values in A-C were statistically compared between dTAGV-1/MZ1 and DMSO with paired t-tests and BH-corrected. Adjusted p-values <0.1 (†) and <0.05 (*) are shown in the corresponding fold-change column (see Supplemental table 1 for statistics and replicate numbers).

We also co-depleted BRD4 and WAPL using MZ1 and dTAGV-1 treatment in the WAPL-dTAG cell line (Fig. 5B). If BRD4 operates through NIPBL and loop extrusion, WAPL co-depletion should (at least partially) rescue this [56,95]. However, co-depletion of WAPL and BRD4 further exacerbated the decreased activation for activator pools at 96 and 409kb, in contrast to what would be expected if BRD4 was operating via loop extrusion. MZ1 reduced the increase in activation from the 848kb activator pool we had seen upon WAPL depletion, but this is also consistent with MZ1 alone slightly decreasing activation from this location and could therefore be independent of loop extrusion. Overall, these results suggest that BRD4 depletion affects gene activation away from the promoter at both short- and long-range, and that loop extrusion and BRD4 affect enhancer function through parallel pathways.

Besides affecting median expression, perturbation can change cell-to-cell variability, also known as transcriptional noise. To test this, we measured the coefficient of variation (CV) of reporter expression. WAPL and NIPBL depletion had no effect on CV when activators were targeted at the *Shh* promoter, but BRD4 depletion was associated with increased CV at the promoter, even though median expression across the cell population was not significantly altered (Fig. 5C). BRD4 depletion also resulted in increased CV from all other activator sites tested. WAPL depletion led to minor changes in CV from non-promoter targeted sites, with variable directions of change. In contrast, NIPBL depletion decreased CV from 96kb (with unaltered median transcription), 206kb, and 848kb (Fig. 5C). These changes were not mirrored in the CV of EBFP2, transcribed from the transfected plasmid, suggesting this is not due to global changes affecting, for example, post-transcriptional regulation (Suppl. Fig. 7C). Therefore, BRD4 appears to be a general suppressor of variability, likely due to its role in regulating transcriptional bursting [96,97]. Our data suggest that loop extrusion tends to increase variability when activation occurs from distal locations.

### Activation changes correlate with changes in chromatin contacts

To test the relationship between chromatin contacts and spatial distances (between activation site and pShh) with distal gene activation, we correlated the different types of data. Micro-C contacts measured by RCMC at 10kb resolution were correlated to smaller DNA-FISH median distances and larger fraction of alleles <350 nm (Suppl. Fig. 7D). As the level of activation using different pools cannot be directly compared, we performed separate correlations for each pool. Activation from 409kb and 848kb correlated with distance, proximal alleles, and contacts. Activation from the closest enhancer (SBE6, 96kb) correlated with RCMC but not with DNA-FISH (Suppl. Fig. 7D).

To expand the number of data points and to compare across gRNA pools, we compared the fold change in activation potential with the fold change in chromatin contacts by RCMC upon treatment. These data include additional measurements (the 206kb pool and dose titrations) than above and showed a clear correlation between contact and activity change (Fig. 5D). To make the same comparison for *Sox9* activation, we took advantage of published NIPBL and WAPL depletion micro-C data at lower resolution (20kb) as proxies for changed chromatin contacts in our cell lines [39,57]. Also here, the change in chromatin contacts correlated with the change in activation (Fig. 5E). With these data we cannot deduce the quantitative relationship (e.g. log-linearity) between contacts and activation, but the correlations are in agreement with perturbation of cohesin co-factors mediating the effect on distal activation by altering chromatin contacts.

### Cohesin co-factor perturbation affects other enhancer-driven genes

To test the effect of cohesin on gene expression more broadly, we analysed published RNA-seq data from RPE1 cells treated with siRNA against various cohesin co-factors or JQ1, which inhibits BRD4 [43]. These transcriptional changes will include both direct and indirect effects and will be confounded by multiple factors including mRNA half-life. While which genes are regulated by which enhancer is unknown in these cells, we used enhancer-gene link predictions (ENCODE rE2G [11]) and categorised genes as promoter-regulated (no enhancer), or regulated by short-(<10kb), mid-(10-50kb), or long-(50kb-2Mb) range enhancer(s). This workflow is derivative of a previous study which found that cohesin preferentially affects long-range regulated genes [63]. We used differentially regulated genes annotated in the original study and excluded rE2G predictions with ambiguous assignments, leading to a final set of 1,672 genes. First, we compared the magnitude of expression changes (fold-change) of predicted long-range vs short-range regulated genes. siRNA against RAD21, NIPBL, or both, was statistically enriched for downregulation of long-range genes (Fig. 6A-B). This agrees with our results above where reduced loop extrusion particularly affects distal enhancers. Second, we compared whether significantly dysregulated genes were unevenly distributed across the different categories of assigned enhancers (Fig. 6C-D). This was true for all perturbations, with RAD21 and MAU2 perturbations most enriched for downregulated long-range vs short-range genes. In contrast, WAPL/PDS5A/PDS5B depletions, which all lead to increased loop length and stalling [39,40,50], were enriched for long-range genes in their upregulated set (Fig. 6C-D). This is in line with our finding of increased cohesin stalling leading to increased distal activation (Figs. 3, 4). Third, we compared the fraction of dysregulated genes controlled by long-range enhancers, or the corresponding enhancers, that were within 5kb of a CTCF peak (Fig. 6E-F). Only WAPL and PDS5A/B upregulated genes were statistically enriched in CTCF binding near upregulated versus downregulated genes and enhancers. While these global analyses cannot directly attribute causality, they are consistent with i) reduced loop extrusion (via RAD21/NIPBL/MAU2 perturbation) leading to downregulation of long-range regulated genes and ii) increased long-range loops (via WAPL/PDS5A/B perturbation) leading to up/downregulation at long-range regulated genes, with those without CTCF near the gene or enhancer being preferentially downregulated and vice versa.

**Figure 6.**
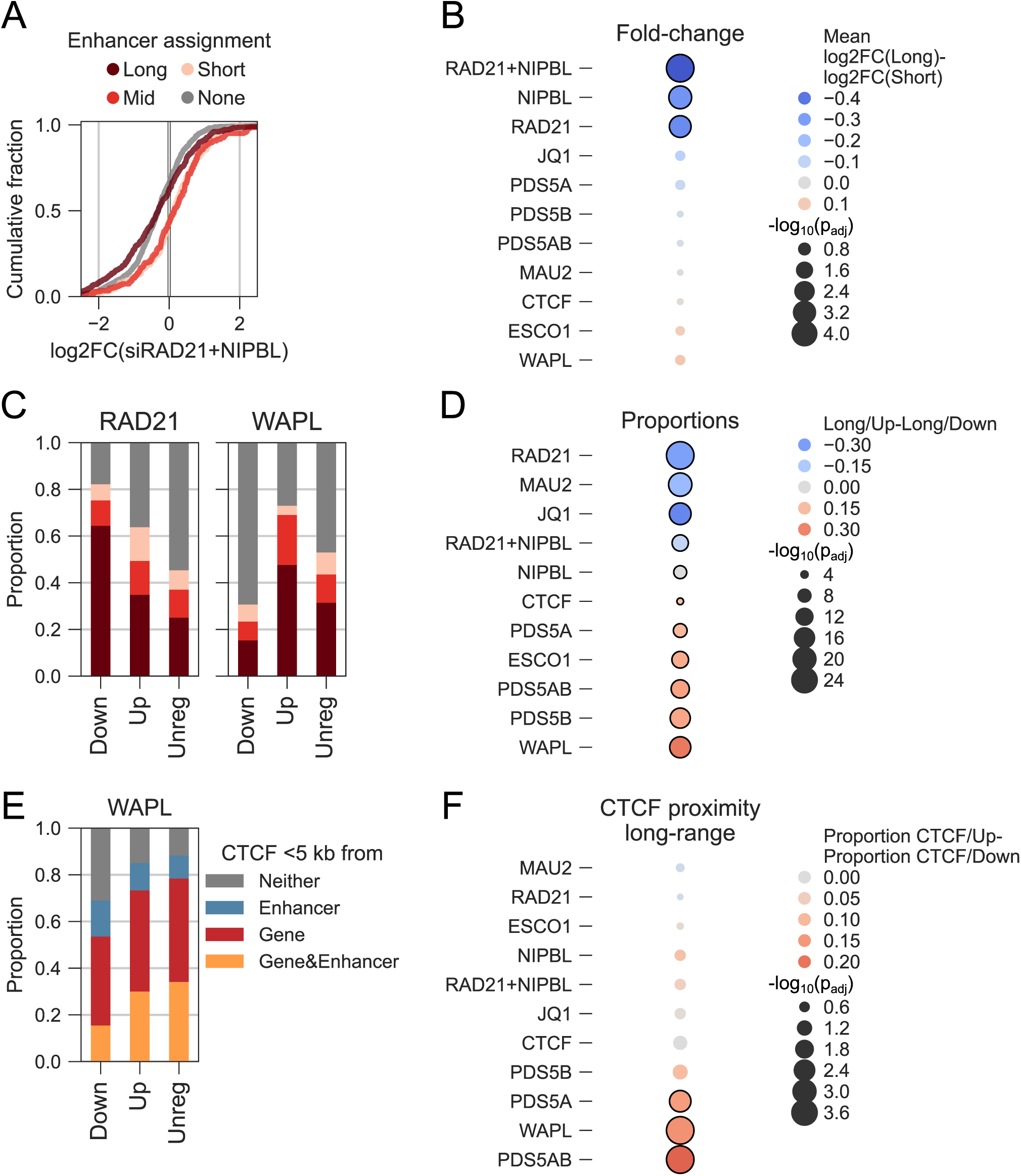
Cohesin co-factor depletion in RPE1 cells preferentially affects long-range enhancer regulated genes. A) Distribution of log_2_ fold-changes of the different gene categories upon treatment with siRNA against RAD21 and NIPBL. B) Heatmap of log_2_ fold-changes (colours) and log-transformed and inverted adjusted p-values (sizes) for the different treatments based on t-tests. Conditions with p_adj_<0.05 have a black outline. C) Distribution of gene categories in the Down-/Up-/Unregulated gene sets for cells treated with siRNA against RAD21 or WAPL. D) Heatmap of differences of proportions of long-range and upregulated genes versus long-range and downregulated genes (Long/Up minus Long/Down, colours) and log-transformed and inverted adjusted p-values (sizes) for the different treatments based on chi-squared tests. Conditions with p_adj_<0.05 have a black outline. E) Distribution of CTCF proximity categories in the long-range and Down-/Up- /Unregulated set for cells treated with siRNA against WAPL. F) Heatmap of differences of proportions of long-range CTCF-proximal genes/enhancers in the upregulated versus downregulated gene sets (CTCF/Up minus CTCF/Down, colours) and log-transformed and inverted adjusted p-values (sizes) for the different treatments based on chi-squared tests. Conditions with p_adj_<0.05 have a black outline. Data in these panels come from Nakato et al. 2023 [43] (see Supplemental table 1 for statistics).

We also found that treatment with the JQ1 BET inhibitor (of BRD4) led to preferential downregulation of long-range regulated genes (Fig. 6D, Suppl. Fig. 7E). The study from which we used the data found minimal changes in chromatin structure upon JQ1 treatment [43]. Together with previous work [63,98], and our results using MZ1 (Fig. 5A-B), these results suggest BRD4 is particularly required for long-range regulated genes albeit not via an impact on cohesin-mediated loop extrusion.

### Dual activation follows a non-linear response curve

Non-additive and context-dependent effects on transcription have recently been demonstrated when combining or perturbing multiple different enhancers [4,5,8,11]. This could arise from synergy between the different TFs and co-activators recruited to different enhancers. Whether non-additive effects occur when the same activators are targeted simultaneously at different positions, and at proximal vs distal positions, has not been examined. To test this in our minimal context, using only one type of TF activator, we transfected a constant dCas9-VPR amount with a half dose of a given gRNA (or x100 gRNA pool) together with either a half dose of a non-targeting guide (single activation) or another gRNA (or x100 gRNA pool; dual activation) (Fig. 7A). We tested activation from multiple pairs of distal and promoter-proximal (pShh-1 and pShh-2) sites. To see if saturation could be reached, we also included a promoter-proximal x100 gRNA pool (pShh x100), which led to massive upregulation of the reporter (Suppl. Fig. 8A). In all cases, dual activation led to higher reporter expression than single activation (Fig. 7A).

**Figure 7.**
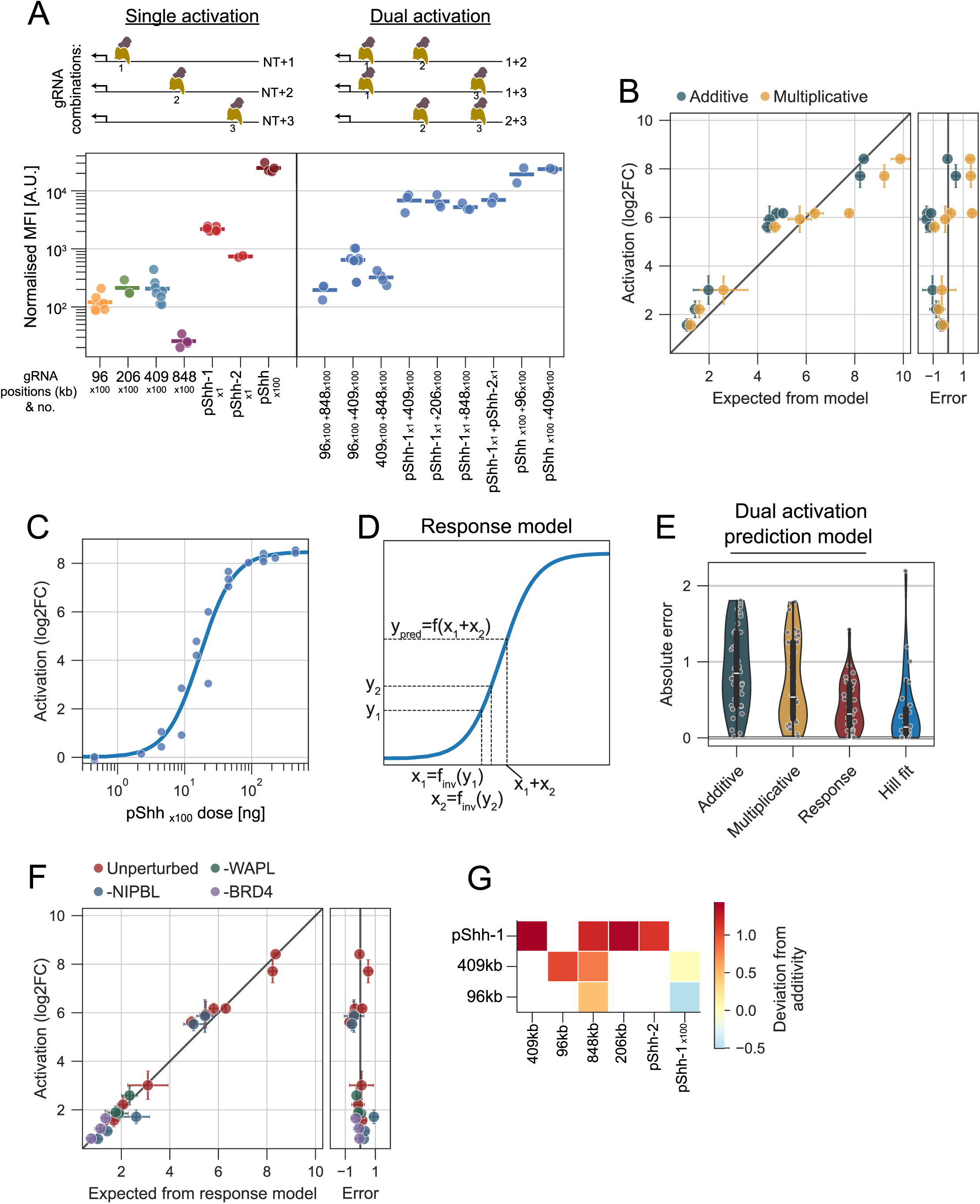
Cooperative activation follows a non-linear response function independent of distance and perturbations. A) Top: Schematic of single and dual activation using different gRNA pool combinations. NT is a non-targeting gRNA. Bottom: Mean mGreenLantern fluorescence (MFI), normalised by subtracting the value of NT-transfected cells for Shh::mGL reporter cells transfected with dCas9-VPR and a combination of different gRNA or gRNA pools (x axis) together with an NT guide (single activation) or with another gRNA or gRNA pool (dual activation). B) Estimation of dual activation based on single activation levels using an additive and multiplicative model. Measured log_2_ fold-change (MFI_sample_/MFI_background_) are shown on the y axis. On the left x axis, the expected dual activation values for the models are shown. On the right axis, the deviation from the measured value (error) is shown. Error bars represent standard deviation. C) MFI for mGreenLantern fluorescence normalised to NT-transfected cells by division (points, log_2_ fold-change; y axis) for Shh::mGL reporter cells transfected with dCas9-VPR and different doses [ng] of a gRNA pool targeting the *Shh* promoter (x axis). A Hill function was fitted to the data (line). D) Schematic of the response model. Two single activation values y1 and y2 and the inverse of the fitted functions are used to find the corresponding x_1_ and x_2_ values and the dual activation output predicted using the fitted function f(x_1_+x_2_). E) Violin show the absolute error (deviation from measured values) of the additive, multiplicative, and response model to predict dual activation, as well as for the Hill fit in panel C. Boxes show interquartile range (Q1-Q3) and whiskers show remaining data range. F) Estimation of dual activation based on single activation levels using the response model as described in panel C. Measured log_2_ fold-change (MFI_sample_/MFI_background_) are shown on the y axis. On the left x axis, the expected dual activation values for the model is shown. On the right axis, the deviation from the measured value (error) is shown. The points are coloured based on the perturbation of NIPBL or WAPL using dTAGV-1 in the respective cell lines, or BRD4 using MZ1. Error bars represent standard deviation. G) Heatmap of the average deviation from an additive model shown for pairs of guides in the unperturbed state, where values >0 represent super-additivity.

To test if dual activation combined in a predictable way, we compared the observed values to additive and multiplicative models of single activation (Fig. 7B). While the additive model tended to underestimate activation, the multiplicative model better predicted the combined response. However, at higher values the multiplicative model led to overestimation, suggesting saturation is being reached. As neither of these models captured the range of experimental values, we wondered whether the combinatorial activation could be described by a non-linear dose-response curve of the dCas9-VPR activator. Indeed, titrating the concentration of the proximal x100 gRNA pool, yielded a typical sigmoidal response pattern as is often seen for TFs (Fig. 7C) [99]. We fitted multiple sigmoidal functions, of which a Hill function best described the data (Suppl. Fig. 8B-C). To apply this as a model for the combinatorial situation, we took the observed value for single activation and found the corresponding x axis value of the curve using an inverse Hill function, i.e. the equivalent dose of proximal x100 gRNA pool that leads to that level of activation. We then calculated the value of the forward Hill function for the combined x axis values, which we call the response model (Fig. 7D). The response model captured the range of combinatorial values better than both the additive and multiplicative models, with absolute errors slightly higher on average than those of the fitted data itself (Fig. 7E). Note that the data of the fitted curve are unrelated to the single/dual activation data and includes only one of the same guide pools. The prediction of the response model is therefore not a simple result of data fitting.

The ability of the response model to predict dual activation implies that signals combine independently of the location of activation, as the model was derived only with proximal guides. Indeed, the prediction error was similarly low for pairs of distal sites, pairs of proximal sites, and combinations of proximal and distal sites (Suppl. Fig. 8D). This strongly suggests that proximal and distal activation are equivalent in nature, but that proximal activation is simply more efficient. As we have shown that perturbing loop extrusion can have a large influence on distal activation, we wanted to see if this would break the predictive ability of the response model when using combinations of distal or proximal and distal guides to target the activators. In contrast, the error was no greater for dual activation in the presence of perturbed NIPBL or WAPL (Fig. 7F, Suppl. Fig. 8E). We also performed single and dual activation upon BRD4 depletion using MZ1, which also did not break the ability of the response model to predict dual activation (Fig. 7F, Suppl. Fig. 8E).

Overall, the response model predicted dual activation for many combinations of activation sites and perturbations (Fig. 7F). Notably, other sigmoidal fits also performed better than the multiplicative and additive models on the totality of the data, and better than a multiplicative model with a saturation point set at the highest observed response value (capped multiplicative; Suppl. Fig. 8F-G). The non-linearity of the response curve means that a given signal can lead to a range of responses when compared to an additive model, depending on the other signal it is paired with, which could be interpreted as context-dependent cooperativity (Fig. 7G). We conclude that, at least in the context studied here, there is an equivalence of proximal and distal gene activation. Furthermore, combinatorial activation follows a non-linear response curve, independently of perturbations of BRD4 or loop extrusion.

## Discussion

### Cohesin co-factor perturbation affects long-range gene activation

We describe, to our knowledge, the longest-range synthetic gene activation to date (up to 1.2Mb) achieved by recruiting large numbers of dCas9 proteins fused to a strong synthetic activation domain (VPR) to sites across the *Shh* and *Sox9* TADs. Using this CRISPRa activation system, we found that depletion of the cohesin co-factor NIPBL, required for loop extrusion, led to reduced transcription when activation was driven from distal regions (>100kb). Regions further away were more affected and the effect was NIPBL dose-dependent. Depletion of WAPL, which drives cohesin unloading and turnover, affected distal gene activation dependent on the proximity to CTCF sites. WAPL depletion increased activation from regions where cohesin was stalled, i.e. near CTCF binding sites, while it decreased activation from regions without CTCF. We substantiated these findings by reanalysing published global transcriptional measurements upon cohesin co-factor knockdown in RPE1 cells and computational enhancer assignments. It is well-established that cohesin stalling by WAPL depletion is CTCF position-dependent [50,51,86], but we do not exclude that other factors also play a role. Together with other studies (for example [61–63,65]), the data presented here strongly supports a model of cohesin-driven loop extrusion as an indirect regulator of distally activated transcription.

The changes in reporter activation in different conditions broadly correlate with changes in chromatin contacts between *Shh* and the region where the synthetic activators are bound, suggesting these measured contacts are correlated with functional contacts. However, our data are not sufficient to determine the precise relationship between contacts and transcription at the *Shh* locus. Interestingly, we found that cohesin co-factor perturbation can have unintuitive outcomes. For example, activation from 96kb upstream of *Shh*, devoid of nearby CTCF sites, is insensitive to NIPBL or RAD21 depletion but sensitive to WAPL depletion, i.e. not requiring loop extrusion but affected by loop extrusion. This suggests that the dynamic unloading of cohesin can contribute to distal gene regulation independent of CTCF near the activation site. Besides affecting median expression, we found that loop extrusion drove increased cell-to-cell variability when activation occurred from several distal enhancers. This could be due to increased heterogeneity in chromatin contacts in the presence of loop extrusion, as suggested by modelling [100]. We note that another study found that RAD21 depletion increased expression variability [101].

If loop extrusion impacts distal gene activation, why do several previous studies show minor effects of perturbing cohesin on gene expression? It has been suggested that this could be explained by cohesin being mainly important for gene expression changes but not for steady-state expression [93,102]. However, other work [61,65], and our re-analysis of cohesin perturbations in RPE1 cells, show acute reduction of enhancer-driven gene transcription upon cohesin perturbation. We suggest that the reported minimal effects of cohesin perturbation are instead likely explained by only a minority of genes being mainly dependent on long-range enhancers. This was recently shown for *Sox2*, where the dependence on loop extrusion was only revealed after deletion of a proximal enhancer element [65]. Furthermore, the effect of cohesin perturbation on chromatin structure is variable throughout the genome and affected by factors including, but not limited to, the strength and relative position of CTCF sites and TAD boundaries. For example, while activation of *Shh* and *Sox9* appears cohesin-insensitive up to ∼100kb, *Car2* is sensitive already at ∼20kb [65]. Globally, the variable expression level of cohesin co-factors and cell cycle length/distribution between cell types may also alter sensitivity to perturbation. For example, while we found no effect of partial WAPL depletion in our experiments, WAPL levels have been shown to be important in other contexts such as regulation of the Protocadherin cluster or V(D)J recombination where reduced WAPL favours long-range interactions [103,104]. Importantly, both genes examined here (*Shh* and *Sox9*) reside at the edge of TADs, and CTCF deletion has been shown to be able to affect their transcription levels [105,106]. How having genes more centrally positioned in TADs, and devoid of nearby CTCF sites, affects sensitivity to cohesin will require further studies. We also have not tested the impact of cohesin-independent sources of chromatin contacts, such as affinity between active regulatory elements, which are less easily manipulable [80,107,108].

### Disease-associated perturbations affecting long-range gene activation

By titrating the level of NIPBL depletion, we showed that there is a dosage effect on distal activation. Our data suggest that in the developmental disorder CdLS, where levels of *NIPBL* mRNA are reduced to about 60% of wild-type, there may be a significant problem in the ability of enhancers that are ∼200kb or more from their target genes to function correctly in development. Knockdown of RAD21, SMC1A and SMC3, whose mutation also cause CdLS, also affects distal activation [44,61,63]. CdLS-like syndromes can result from mutations in genes involved in transcriptional regulation including BRD4 [89]. Though no unified mechanism is known, our data are inconsistent with BRD4 operating directly via NIPBL/cohesin, and there was no clear change in chromatin contacts upon BRD4 perturbation. Importantly, BRD4 degradation and JQ1 treatment both lead to preferential downregulation of long-range controlled genes [63], suggesting that CdLS-like disorders might be linked through perturbed distal enhancer function by parallel pathways.

### Synthetic activation reveals principles of distal activation and cooperativity

We used our synthetic activation system to drive *Shh* reporter expression from two locations simultaneously, mimicking activation from multiple enhancers, but notably using the same activator, and therefore presumably recruiting the same co-activators, at both sites. We found that neither an additive nor a multiplicative model, based on the single activators, could capture the range of dual activation outputs. Instead, we found that the combinatorial activation could be predicted by a non-linear response curve determined by titrating activators to the promoter. A plausible conclusion is that distal gene activation is not inherently different in nature to proximal activation, simply much less efficient. Our ability to activate the silent Polycomb-repressed *Shh* gene via activator recruitment at distances of several hundred kb also suggests that distal enhancers do not require proximal enhancers to achieve gene activation, contrary to what has been suggested [109]. However, cooperativity can lead long-range enhancers to have a much larger transcriptional impact in conjunction with other enhancers.

We propose that, at least in the context studied here, a given activation signal has the same impact on the gene regardless of its origin (proximal/distal) or perturbations to WAPL, NIPBL, or BRD4. When combined with another signal, the non-linear nature of the response curve leads to variability in the observed effect on gene expression. This is similar to the concept of dose additivity (Loewe additivity) in pharmacology, where the input signals rather than output responses combine additively [110]. Here, the non-linear Hill function was positively cooperative (n_Hill_=1.73), and hence we observed mainly super-additive effects until at saturation, although the extent of super-additivity varied depending on which two signals were combined. Previous work showed that expression data from partial super-enhancer deletions was best fitted to non-linear models, rather than additive or multiplicative models, suggesting this applies more generally [111]. It will be interesting to test if a non-linear response model can explain other types of cooperativity documented in the literature.

What can explain position-agnostic cooperativity in our assays? Local models involving direct or indirect TF-TF cooperativity or allosteric co-factor bridging do not make sense when the binding occurs hundreds of kb apart. We can also exclude different enhancers recruiting different co-activators that control different steps of transcription, since we assume that the single activator species (dCas9-VPR) recruits the same co-activators at each targeting site. Instead, cooperativity might arise from how the VPR activator impacts the promoter, e.g. through direct recruitment of RNA polymerase II, or by affecting various steps of the transcription cycle [112,113]. While the response model explained our dual activation data well, we cannot exclude that some combinations in certain conditions with larger errors represent genuine outliers from the model where additional mechanisms operate.

### Model of enhancer activation based on contact and response curves

A simple model of enhancer activation can be envisioned given our data and results from previous studies (Fig. 8). First, for any given enhancer or activator, the resulting transcription scales with chromatin contacts [24,114]. When placed next to the gene, the enhancer has its maximal influence, and when moved further away contacts will decrease in a locus-specific manner [24,70,72,114]. These contacts are highly affected by loop extrusion dynamics and CTCF. Second, combinations of multiple enhancers or activator signals will combine according to the response model. This can give rise to cooperativity, either positive (synergy) or negative (redundancy) depending on the shape of the curve and the magnitude of the inputs. If one could determine the contact-activation and input-activation curves for a given gene and activator, one could predict any combination of enhancers. This does not explain how or which chromatin contacts lead to activation or which parameters affect the shape of the curves. Furthermore, such a prediction may only be possible with a single type of enhancer or activator. In more complex and biologically applicable scenarios with different and varying TFs per enhancer, there might not be a single response curve, but it might change shape in combination with other signals [115]. For example, cooperative processes can be imagined where certain regulatory elements drive contacts or where different enhancers affect different parts of the transcription cycle [12,112]. It is thus possible that distal and proximal activation are differently affected by specific co-factors and their combinations. How genes integrate signals from distal and proximal regulatory elements with diverse TF repertoires will be an important area of future study.

**Figure 8.**
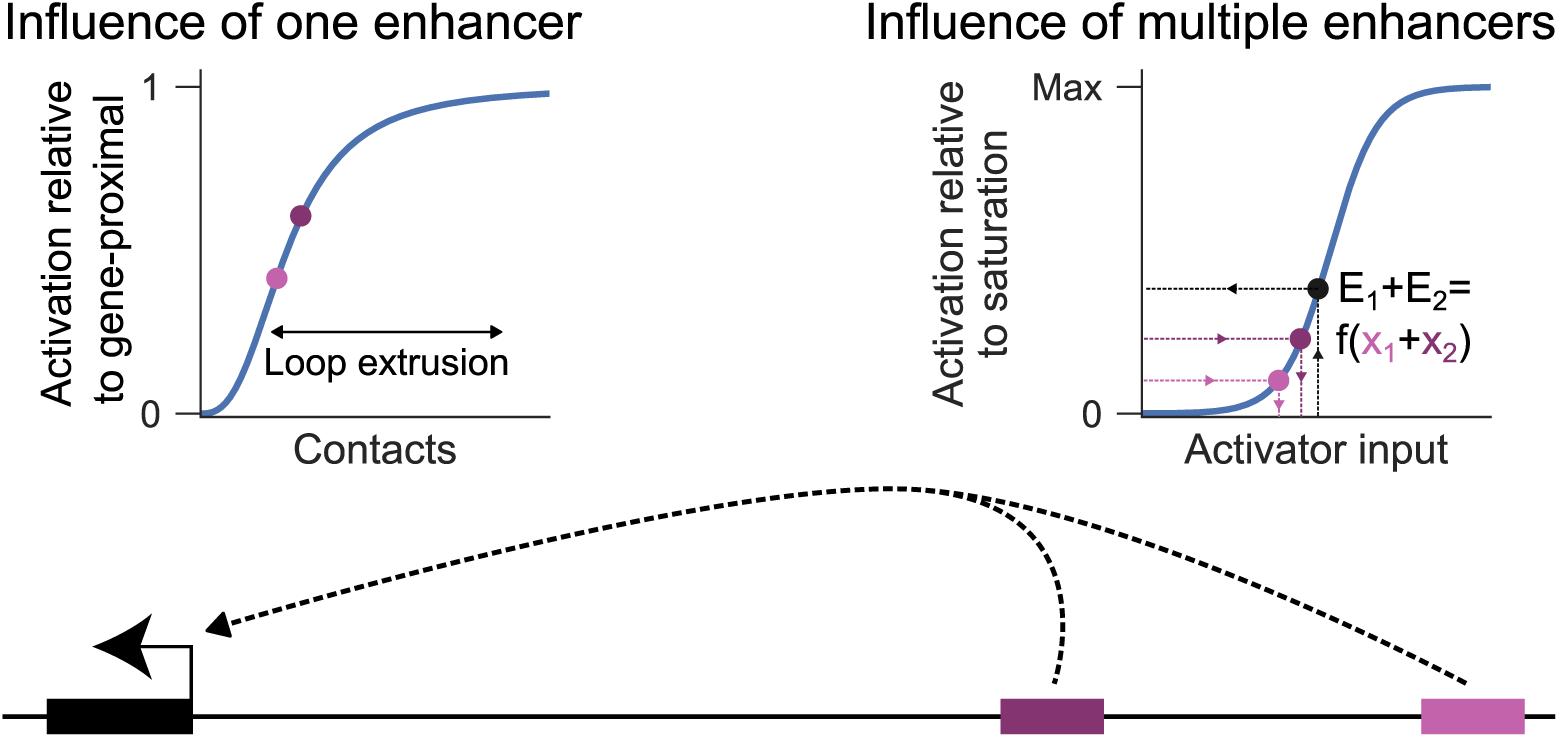
Model of enhancer activation based on contact and response curves. Model depicting how two non-linear relationships affect enhancer-driven transcription. Left: A contact-activation curve shows how distal enhancers have lower influence the less they contact the promoter (see e.g. [24,70,114]). This depends greatly on genomic distance, and the activity of loop extrusion. Right: An input-response curve shows how increasing inputs (number of enhancers/activators) leads to increasing outputs. The combined activity of two enhancers corresponds to their combined individual input signals, rather than their combined individual outputs.

## Materials and Methods

### Cell culture

E14Tg2a-derived mESCs were grown at 37°C, 5% CO_2_ in GMEM BHK21 (Gibco, 21710025) supplemented with 15% fetal calf serum (FCS), 2 mM L-glutamine, 1 mM sodium pyruvate (Sigma, S8636), 1X non-essential amino acids (Gibco, 11140035), 50 mM 2-β-mercaptoethanol (Gibco, 31350-010),1000 U/ml LIF (Merck, ESG1107), and penicillin/streptomycin on 0.1% gelatin-coated tissue culture plastics. For *Sox9* reporter experiments, 3 μM CHIR99021 (A3011-APE, Stratech) and 1 μM PD0325901 (A3013-APE, Stratch) was added to the culture medium. Cells were washed with PBS, and lifted with 0.05% Trypsin-EDTA (Gibco, 25300054) to split every 2 - 3 days, except after clonally sorting into 96-well plates where medium was replaced every two to three days until there were visible colonies. Cell cultures were routinely tested for Mycoplasma and were negative. The RAD21-AID (SCC1-AID) cell line was generously gifted by the lab of Rob Klose [78].

### Transfection

5 x 10^4^ -2 x 10^5^ cells in 0.5 ml medium in 24-well plate wells were transfected immediately after seeding with 500 ng circular plasmid DNA, 1.5 μl Lipofectamine 3000 and 0.5 μl P3000 reagents (Thermo Fisher, L3000015) in 50 μl OptiMEM (Gibco, 31985062). All plasmids were prepared by QIAprep Spin Miniprep (Qiagen, 27104) or Plasmid Plus Midi (Qiagen, 12943). The cell culture medium was changed the next day. For larger culture dishes, cell numbers and reagents were scaled according to tissue culture area.

### Plasmid construction

A list of plasmids used and their origin can be found in Suppl. table 2, and primers used for cloning in Suppl. table 3. A tShh-VP64-mCherry plasmid was generated by amplification of T2A-mCherry, digestion by NheI/BsrGI and ligation into tShh-VP64-EGFP [31]. The pCAG-dCas9-VPR plasmid was generated by amplification of dCas9-VPR from Addgene #175572, digestion with NotI/NheI and ligation into tShh-VP64-mCherry. Homology arms for the *Shh* and *Sox9* reporters were ordered as cloned plasmids from GeneArt (Thermo Fisher). The dTAG-NIPBL and WAPL-dTAG repair templates were modified from Addgene #163882 and #163873 [116] by amplification of HaloTag and restriction/ligation using BsiWI/BsrGI.

The pU6sgEnhBFP plasmid was constructed by ordering a U6-sgEnhBFP construct containing a U6 promoter followed by BbsI (BpiI) cloning sites (based on pX330 [117]) but with a modified sgRNA based on the F+E extension in [118] from GeneArt. This was amplified and cloned into Addgene #49016 [119] using BamHI/EcoRI restriction/ligation. gRNA oligos were annealed and cloned into the pX330-based vectors according to established protocols (see Suppl. table 3) [117]. gRNA pools (Suppl. table 4) were ordered as pools of single oligos (oPools, IDT), amplified, digested with BbsI, and ligated into the pSPgRNA (Addgene #47108 [120]; Fig. 1B, SBE2 pool) or pU6sgEnhBFP plasmid (see Supplementary protocol for details). Note that we observed similar activity using either backbone. All plasmids were verified by Sanger or whole-plasmid sequencing (Plasmidsaurus). gRNA pools were verified by scanning whole-plasmid raw sequencing reads for the gRNA cloning site and mapping the corresponding gRNA entry to the oligo pool. gRNA design and off-target screening was done based on CRISPOR outputs [121].

### Cell line generation

For knock-ins, E14 cells were transfected at a 3:1 molecular weight ratio of repair template plasmid:Cas9/gRNA plasmid. Following selection/sorting and clonal expansion procedures, genomic DNA was harvested using DirectPCR (Viagen, 301-C), and genotyping performed with GoTaq (Promega, M7122) using primers in Suppl. Table 3. For the Shh::mGL reporter, transfection was performed in E14 WT cells and 1 μg/ml Puromycin was used for 4 days to select against untransfected cells. Cells were maintained for a further 11 days to exclude transient fluorescence, then population-sorted for mCherry-positive cells expressed from a PGK promoter flanked by loxP sites (Suppl. Fig. 1A). This population was later transfected with a plasmid transiently expressing Cre recombinase and clonally sorted for mCherry-negative cells. For dTAG-NIPBL cells, the selected clone of Shh::mGL was transfected, and 2 days later treated with 8 μg/ml Blasticidin for 4 days before HaloTag-JF646-positive cells were clonally sorted. For WAPL-dTAG cells, the selected clone of Shh::mGL was transfected and 2 days later HaloTag-JF646-positive cells were clonally sorted. dTAG-NIPBL and WAPL-dTAG clones were karyotyped and confirmed diploid. For the ZRS KO, dTAG-NIPBL cells were transfected with dual gRNAs cloned into pX458 and 2 days later GFP-positive cells were clonally sorted. A negative control clone was derived from similarly treated and sorted cells transfected with another unrelated gRNA pair, and confirmed unedited. For the *Sox9* reporters, dTAG-NIPBL or WAPL-dTAG cells were transfected and after one day treated with 1.5 μg/ml Puromycin for 1 day, and 5 days later transfected with a plasmid expressing dCas9-VPR (Addgene #63798 [122]) and a cocktail of gRNAs targeting the *Sox9* TAD, before sorting cell populations as mScarlet3-positive.

### Small molecule treatments

dTAGV-1 (Tocris, 6914) and MZ1 (BPS Bioscience, BPS-82056) were resuspended in DMSO, and the equivalent concentration DMSO used as control. Following seeding, treatments were immediately added to cells except for RCMC experiments where treatments were added one day after seeding. dTAGV-1 was used at 250 nM for all experiments except where specified, and MZ1 was used at 100 nM. Auxin (Merck, I5148) was dissolved in water and used at a final concentration of 500 μM for 6 hours, with untreated cells as control. For 48 hour experiments, medium and treatments were refreshed after ∼24 hours.

### Activation experiments

Cells were transfected with either a combined gRNA/dCas9-VPR plasmid (Addgene #175572) or an equal molecular weight of a pCAG-dCas9-VPR and gRNA expressing plasmid (250 ng of each) immediately after seeding cells and treatment with DMSO/dTAGV-1/MZ1 where specified. The combined expression vector (Addgene #175572) or pCAG-dCas9-VPR together with pSPgRNA (Addgene #47108) was used in Fig. 1B and Suppl. Fig. 2A-D, and mCherry-positive cells analysed. All other *Shh* activation experiments was done with a combination of pCAG-dCas9-VPR and pU6sgEnhBFP, and mCherry/EBFP2 double-positive cells analysed. For single/dual activation experiments, to avoid gRNA dose being limiting, 50 ng pCAG-dCas9-VPR and 450 ng pU6sgEnhBFP (gRNA) plasmids were used, with 225 ng of one gRNA (or pool) plus 225 ng NT (single activation) or 225 ng of a second gRNA or pool (or pool; dual). For titration of the pShh x100 gRNA pool, cells were transfected with 50 ng of pCAG-dCas9-VPR and different amount of the targeting pool mixed with NT guide to reach a total of 450 ng. For *Sox9* reporter activation, SP-dCas9-VPR (Addgene #63798) without a fluorescence marker was used together with pU6sgEnhBFP.

### Live cell staining

To stain DNA, cells were trypsinised and incubated with 1.6 mM Hoechst33342 (Thermo Fisher, H3570) diluted in culture medium for 15 minutes (min) at 37°C, then spun down and resuspended in cold PBS+1% FCS. To stain for the HaloTag, adherent cells on plates were incubated with 200 nM HaloTag-JF646 (Promega, GA1120) diluted in culture medium for 15 min at 37°C, then washed with fresh unstained culture medium for 5 min at 37°C, then trypsinised and resuspended in cold PBS+1% FCS.

### Flow cytometry, fluorescence-activated cell sorting, and data analysis

Trypsinised cells were kept on ice in PBS+1% FCS prior to flow cytometry. All flow cytometry experiments were performed on a LSR Fortessa X-20 (BD Biosciences) with laser-filter combinations: 355-450/50 (Hoechst33342), 405-450/50 (EBFP2), 488-525/50 (mGreenLantern), 561-610/20 (mCherry/mScarlet3), 640-670/30 (HaloTag-JF646), except for one replicate in Suppl. Fig. 2D which was performed on a Cytoflex S (Beckman Coulter). Sorting was performed on ARIA II (BD Biosciences) and Cytoflex SRT (Beckman Coulter). Gating was performed using FlowJo v10.10, selecting for singlets and where applicable, transfected cells (mCherry+, EBFP2+, or both, depending on the experiment). Single-cell measurements or summary statistics were exported to be further analysed in custom Python notebooks. Samples with <100 measurements were discarded. Normalised MFI values were obtained by subtracting the median fluorescence of the corresponding NT guide transfected population from the median of the sample. For min-max normalisation, a negative control (HaloTag-JF646-stained WT or NT-transfected reporter) was used as the minimum value and the corresponding untreated condition was used as the maximum value. For the *Sox9* reporter, thresholding was achieved by selecting the value of mScarlet3 signal where the average of the NT samples per cell line and replicate was <1% and reporting the percentage above this value.

### Pluripotency assay

Cells were seeded in 6-well plates in normal culture medium and allowed to clonally expand in the presence of DMSO or dTAGV-1. Medium and treatments were refreshed every 2-3 days. On day 7, cells were fixed with 4% paraformaldehyde (PFA; Thermo Fisher, J19943.K2) for 10 min at room temperature (r.t.), washed with PBS, and stained with Vector Red Alkaline Phosphatase substrate kit (Vector Labs, SK-5100) in 100 mM Tris-HCl pH 8.0 in PBS for at least 1 hour at r.t., and washed with PBS.

### Immunofluorescence

dTAG-NIPBL cells were grown on 0.1% gelatinated-coated glass slides in normal culture medium supplemented with 100 nM MZ1 or DMSO for one day and then fixed with 4% PFA, washed with PBS, and permeabilised with PBS with 0.5% Triton X-100. Slides were blocked with PBS with 1% bovine serum albumin (BSA; Merck, 10735086001) for 3 hours at r.t. and incubated with 1:1000 diluted anti-BRD4 antibody (ab128874) in PBS, 1% BSA at 4°C overnight, washed with PBS and incubated with 1:1000 diluted goat anti-rabbit secondary antibody coupled to Alexa Fluor 488 (Thermo Fisher, A11034) in PBS, 1% BSA for 1 hour at r.t. and washed with PBS. Slides were stained with 50 ng/ml 4,6-diaminidino-2-phenylidole (DAPI), mounted in VectaShield (Vector Labs), and sealed with nail varnish.

### DNA-FISH

Cells were grown on 0.1% gelatinated-coated glass slides, fixed in 4%PFA, permeabilised in PBS/0.5% Triton X, dried and frozen at -80°C. Slides were defrosted, incubated in 100 μg/ml RNase A in 2x SSC for 1 hour at 37°C, washed briefly in 2x SSC, put through an alcohol series and air dried. Slides were oven heated to 70°C for 5 min, denatured in 70% formamide/2x SSC (pH 7.5) for 40mins at 80°C, cooled in 70% ethanol on ice for 2 min and dehydrated through 90% and 100% ethanol for 2 min each before air drying. 1μg of each fosmid DNA was labelled by Nick translation to incorporate green-dUTP (Enzo Life Sciences), AlexaFluor 594-dUTP (Invitrogen) or digoxigenin-11-dUTP (Roche). 100ng of each fosmid, 6ul of Cot1 DNA per fosmid, and 5ug of sonicated salmon sperm DNA were ethanol precipitated in a spin-vac and the pellet re-suspended in 30ul of hybridisation mix. Probes were then heated to 80°C for 5 mins to denature and reannealed at 37°C for 15 mins. Probes were hybridised to the slide under a sealed coverslip overnight at 37°C. Slides were washed the next day for 4 x 3 min in 2x SSC at 45°C, 4 x 3 min in 0.1x SSC at 60°C and washed briefly in 4 x SSC/0.1%Tween20. Digoxigenin probes were detected incubating with anti-digoxigenin (Roche) and AlexaFluor 647 (sheep) antibodies for 45 min each and washes of 4 x SSC/0,1%Tween20 inbetween. Slides were finally stained with DAPI at 50ng/ml, mounted in Vectashield (Vector Labs) and sealed with nail varnish.

Fluorescence images were acquired using a Photometrics Prime BSI CMOS camera (Photometrics, Tuscon, AZ) fitted to a Zeiss AxioImager M2 fluorescence microscope with Plan-Apochromat objectives, a Zeiss Colibri 7 LED light source, together with Zeiss filter sets 90 HE, 92 HE, 96 HE, 38 HE and 43 HE (Carl Zeiss UK, Cambridge, UK). Image capture was performed in Zeiss Zen 3.5 software. Images were deconvolved using a calculated point spread function with the constrained iterative algorithm of Zeiss Zen software

### Region Capture Micro-C

Micro-C library preparation was performed according to Goel et al. [80]. Briefly, trypsinised cells were sequentially crosslinked with 3 mM disuccinimidyl glutarate (DSG, ThermoFisher, 20593) in PBS for 35 min at r.t. with gentle mixing, after which formaldehyde (ThermoFisher, 28926) was added to a final concentration of 1%. The double crosslinking reaction was incubated for further 10 min at r.t. with gentle mixing and quenched with Tris-HCl pH 7.5 at a final concentration of 0.375 M for 5 min. Aliquots of 5 x 10^6^ cells were snap frozen in liquid nitrogen and stored at -80°C. Cell pellets were lysed into nuclei in complete MB1 (50 mM NaCl, 10 mM Tris-HCl pH 7.5, 5 mM MgCl_2_, 1 mM CaCl_2_, 0.2 % NP40, protease inhibitor cocktail) at 1 million cells/100 μl for 20 min in ice. Nuclei were spun down at 1750 g for 5 min at 4°C and washed once with complete MB1. Chromatin was in situ fragmented at 37°C for 20 min with MNase (NEB, M0247S) in 100 μl per 1 million cells of MB1. The appropriate amount of MNase was determined by a titration assay with the aim of digesting chromatin to 80-90% mononucleosomes/10-20% dinucleosomes [80]. MNase digestion was stopped with EGTA at a final concentration of 4 mM. Fragment ends were repaired, biotin labelled and proximity ligated as described [80]. After de-crosslinking and DNA extraction, dinucleosome DNA fragments were isolated and extracted form a 1% agarose gel. DNA ends were polished, blunted and ligated DNA was isolated by pulling down biotin-labelled fragments. Illumina library preparation was performed using NEBNext Ultra™ II DNA Library Prep Kit for Illumina (NEB, E7103). Library PCR was performed using index primers (NEBNext Multiplex Oligos for Illumina, Dual Index Primers Set 1; NEB, E7600S) supplemented with 1x EvaGreen (Jena Bioscience, PCR-379). 1/5 of the volume was used to run a test qPCR to determine the appropriate number of cycles by which libraries are amplified to linear phase.

Biotin-bait preparation using fragmented BACs covering the regions of interest was adapted from Golov et al. [123]. 5 μg of pooled equimolar amounts of the selected BACs (Suppl. table 5) were sonicated using MSE Soniprep 150 Plus at medium power for two 30 second pulses on ice to shear DNA to a size of 150-700 bp. Equimolarity was calculated by qPCR using primers against the chloramphenicol resistance gene (Suppl. table 3). Forked adapters (Suppl. table 3) were annealed in NEBuffer 2 at 1X (NEB, B7002S) to a final concentration of 20 µM in a thermal cycler set to heat at 98°C and gradually cool down to 4°C with a 1°C per 20 sec gradient. Shredded BACs were subjected to end-preparation, A-tailing and adapter ligation as described [123]. BAC baits were PCR amplified using 5’ biotinylated forward and reverse primers (Suppl. table 3) in the following reaction mix: 10 µl 2X NEBNext Ultra™ II Q5 Master Mix (NEB, M0544S), 1 µl of each 10 µM bio-primer, 2 µl adapter-ligated BAC DNA, 1X EvaGreen. The number of amplification cycles was determined by the linear phase in a test qPCR. Around 3.5 µg of biotinylated baits were obtained from 25 reactions.

Enrichment of the regions of interest was adapted from Golov et al. [123]. Equal amounts of all libraries corresponding to each sample, and amplified with different barcodes, were pooled and captured as a pool. One hybridization reaction was done per 1 µg of library pool. If more than 4 samples were being captured, more than one hybridization reaction was performed. Hybridization and biotin-pulldown was done as described [123]. Singly enriched 3C libraries were amplified by qPCR using p5/p7 primers (Suppl. table 3) as described above. PCR reactions were performed with the DNA still bound to the streptavidin beads. The number of amplification cycles was determined by the linear phase in a test qPCR. A second hybridization reaction, biotin pulldown and PCR amplification was performed as before. The pool was sequenced using Illumina Nextseq 2000 using a single P2 flow cell (paired-end 50-bp reads).

### Hi-C/Micro-C data analysis

Published datasets used can be found in Suppl. table 6. Visualisation of ICE-normalised chromatin contact maps was done using resgen.io [124]. RCMC data were processed with the distiller-nf v0.3.3 (https://github.com/open2c/distiller-nf) pipeline, with mapq30 filtered and ICE-balanced [125] matrices used. A blacklist containing all regions outside the capture regions was used in the ICE normalisation. Alignments were done to the default mm10 genome (visualisations, P(s) curves) or to a custom mm10 where chr5 is changed so that the *Shh* sequence represents the Shh::mGL reporter sequence (correlations). DMSO/dTAGV-1 contact maps were merged from 2 (WAPL-dTAG) or 3 (dTAG-NIPBL) replicates. dTAG-NIPBL dose titrations were from a single replicate, and fold changes calculated against the paired DMSO (0 nM) condition. Contact values were derived using cooler and cooltools [126,127]. P(s) curves were generated using the cooltools function expected_cis, with a *view_df* excluding chr5, and settings “*smooth=True, aggregate_smoothed=True, clr_weight_name=None*”, and the output normalised to the area under the curve.

### Transcriptomic and E2G analysis

Processed transcriptomic data including reads per sample and differentially regulated genes per condition were downloaded from Nakato et al [43]. Fold change values were derived by differential expression analysis using voom based on TMM-normalised values using design ∼0+condition with Treatment vs Control. E2G predictions were downloaded from ENCODE for various retinal samples and merged (Suppl. Table 6) [11], as there was no prediction specifically for RPE1 cells. These were overlapped with the differentially regulated genes from Nakato et al. and split into categories based on distance to enhancer and E2G prediction scores, essentially as in Tjalsma et al. [63]. Long-range genes were defined as those with a predicted enhancer at ≥50kb and <2Mb with a score ≥0.8. Mid-range genes were defined as those with a predicted enhancer ≥10kb and <50kb with a score of ≥0.8 but no longer-range enhancer at a score of ≥0.3. Short-range genes were defined as those with a predicted enhancer >2kb and <10kb with a score of ≥0.8 but no longer-range enhancer at a score of ≥0.3. No-enhancer genes were defined as those without any predicted enhancer at >2kb with a score of ≥0.3. CTCF peaks were merged with BEDTools [128] from processed files from Nakato et al [43].

### Statistical analysis and graphing

All data were analysed using numpy [129] and pandas [130] data frames in Python [131] jupyter notebooks [132], and graphs generated using matplotlib [133] and seaborn [134]. All statistical tests used were scipy [135] implementations and can be found in Suppl. Table 1. Generally, paired t-tests were used where paired samples were available, otherwise independent t-tests were used. To compare category distributions (Fig. 6D,F), a chi-square test was used, and correlations were tested using Pearson’s correlation. All p-values were adjusted by Benjamini-Hochberg (FDR) correction, implemented in statsmodels [136], for all samples within a graph and p_adj_ values <0.05 considered significant.

### Curve fitting and dual activation models

All curve fitting was done using the curve_fit function in scipy. The min-max normalised curves (Figs. 2D, 3D) were fitted to the decay function

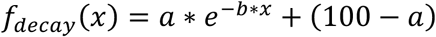

The titrated pShh x100 gRNA dose (x) and log2 fold-change values (y) in Suppl. Fig. 8B were fitted to the following sigmoidal curves:

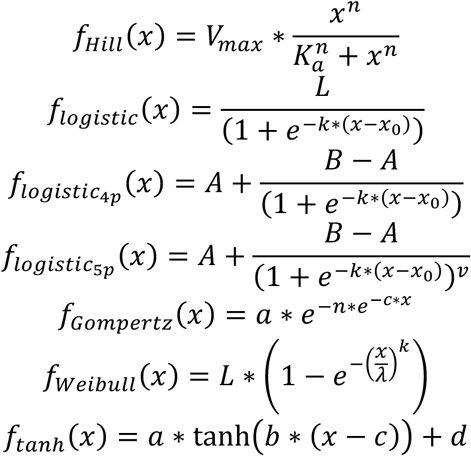

Dual activation was compared to the two corresponding paired single activation experiments performed on the same day. Dual activation was compared to the following additive function:

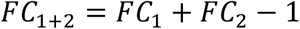

Dual activation was compared to the following multiplicative function:

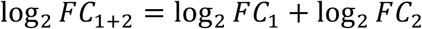

The capped multiplicative was set to cap any values above the mean of the guide with the highest response (pShh x100).

Dual activation was compared to response models using *f*(*x*_A_ + *x*_C_) of sigmoidal functions fitted to the pShh x100 gRNA titration data, where x_1_ and x_2_ were derived from the single activation log_2_FC values using the following inverse functions.

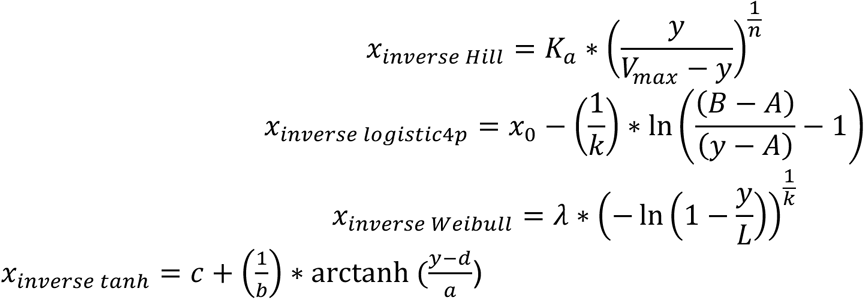

## Supporting information

Supplemental protocol

## Figure legends

**Supplemental Figure 1.**
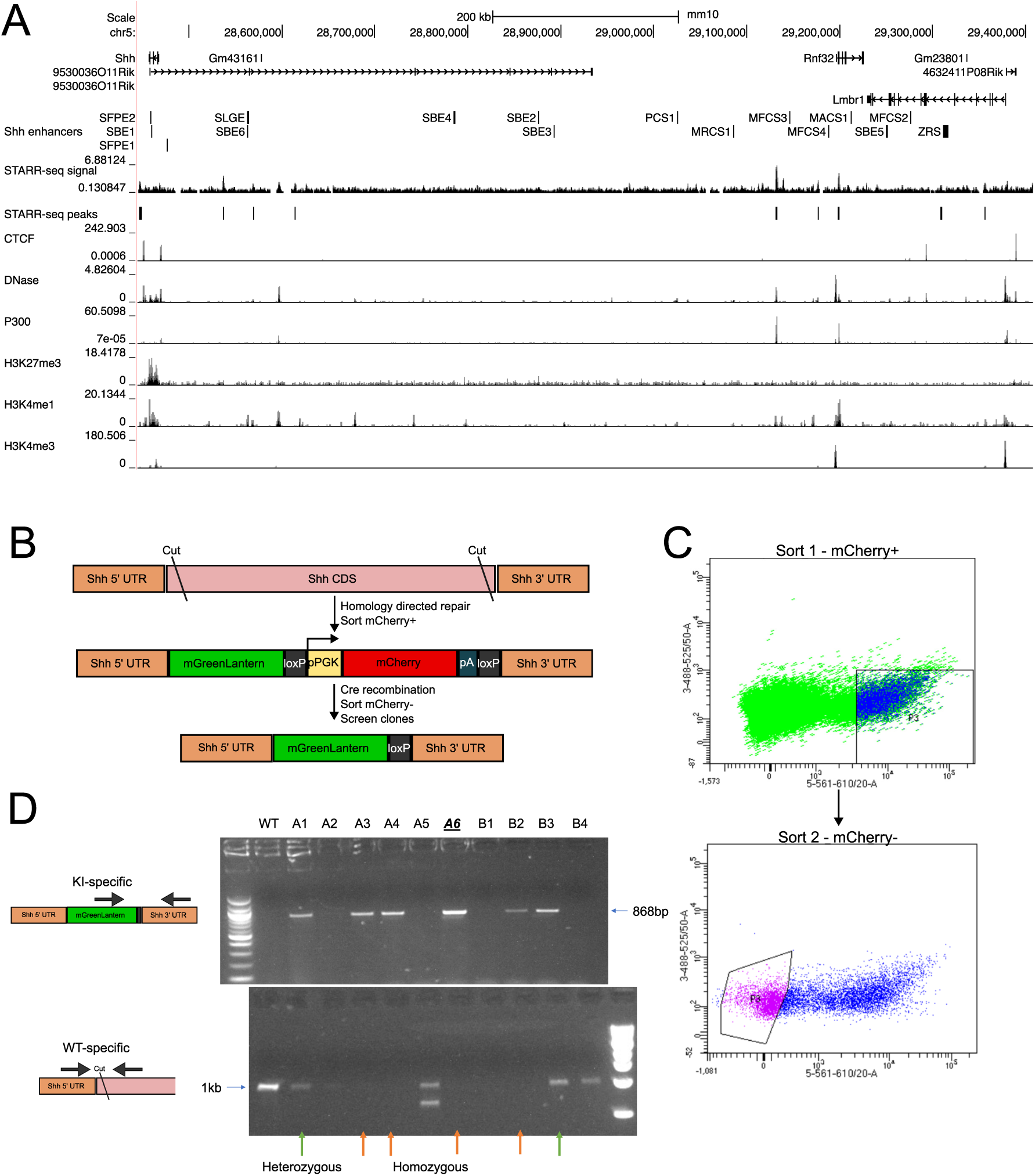
Related to Figure 1. A) UCSC Genome Browser tracks [139] at the *Shh* locus (mm10 genome) together with coordinates for known *Shh* enhancers [16], mESC STARR-seq peaks and signals from [140], and ENCODE mESC tracks for CTCF ChIP-seq, DNase-seq, and H3K27me3, P300, H3K4me1, and H3K4me3 ChIP-seq [141]. B) Schematic of the Shh::mGL reporter generation. Note that the locus has been flipped compared to the genome for clarity. Cas9 was used to cut at the stop and start of the *Shh* coding DNA sequence (CDS), and a repair template containing *Shh* homology arms, mGreenLantern, loxP, PGK promoter (pPGK), mCherry, polyadenylation signal (pA), and loxP was used. After sorting for mCherry-positive cells, Cre recombinase was used to excise PGK-mCherry-pA, and clonal sorting of mCherry-negative cells. C) Fluorescence activated cell sorting (FACS) gates used for the sorting described in panel, with the y axis (green fluorescence) used as a negative signal against mCherry on the x axis. D) Genotyping for Shh::mGL knock-in. Top: Amplification from mGreenLantern to outside the homology arms to verify correct insertion. Bottom: Amplification around *Shh* start codon to exclude WT alleles. The underlined clone A6 was selected for use.

**Supplemental Figure 2.**
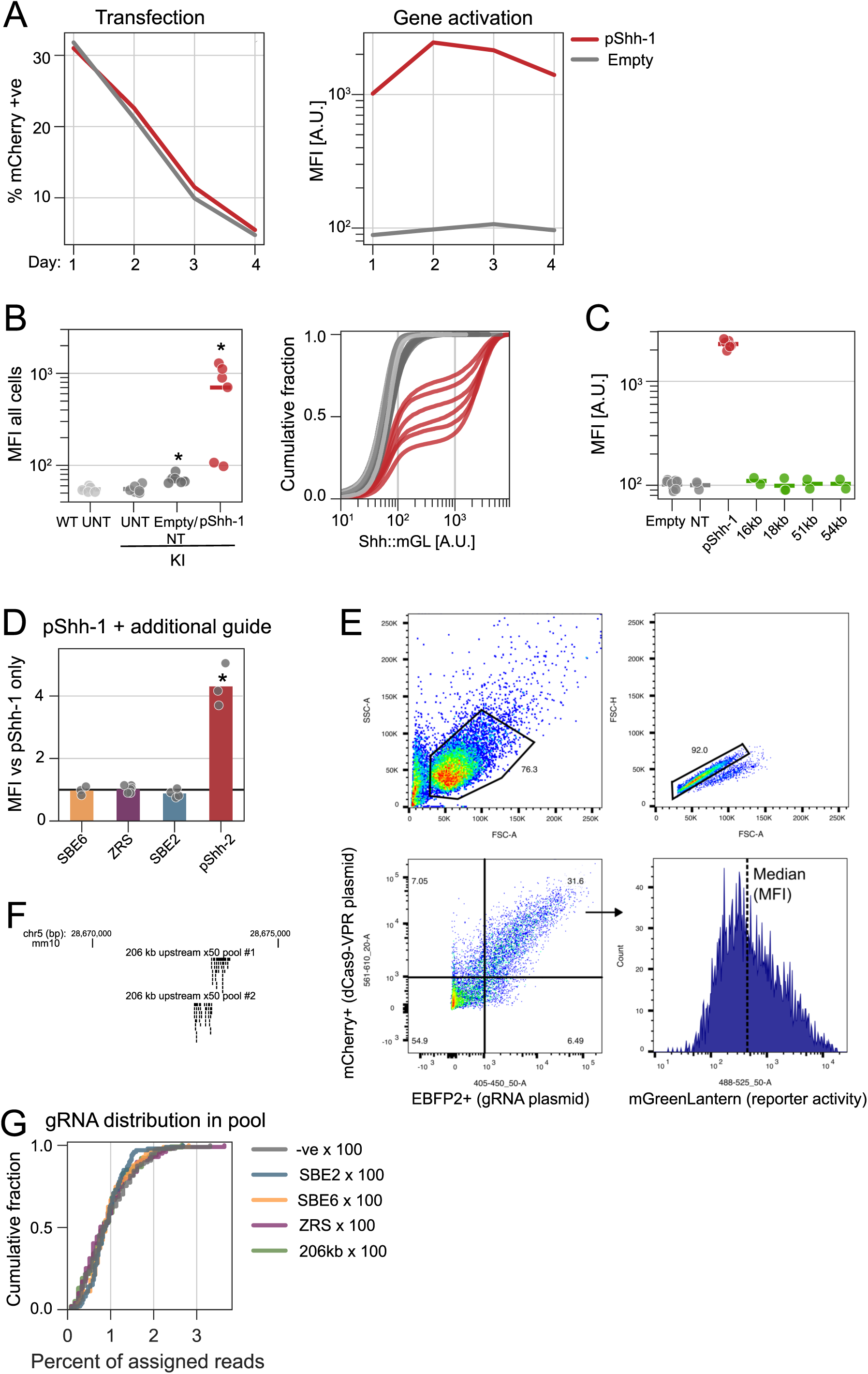
Related to Figure 1. A) Flow cytometry of Shh::mGL cells transfected with dCas9-VPR and an empty guide vector or a guide targeting the promoter (pShh-1) and analysed 1-4 days after transfection for percentage of mCherry-positive cells (expressed from the transfected plasmid, left) and mGreenLantern fluorescence (expressed from the Shh::mGL reporter) of the transfected population (median fluorescence intensity (MFI), right). B) Flow cytometry of wildtype (WT) or Shh::mGL reporter (KI) cells analysed for mGreenLantern fluorescence upon no transfection (UNT), or transfection with dCas9-VPR and no (empty) or a non-targeting guide (NT) or pShh-1. Left: MFI. Right: Cumulative distribution of single-cell fluorescence values used to derive the median, with colour coding as in the left panel, with grey shades representing negative controls and red representing promoter-target activation. Each dot and line is a replicate. C) MFI for Shh::mGL cells transfected with dCas9-VPR and guides targeting to different locations analysed for mGreenLantern fluorescence. Green dots (16kb, 18kb, 51kb, and 54kb) represent single gRNAs targeted at the specified distance upstream of *Shh*, grey dots represent negative controls and red represent promoter-target activation. Empty/NT/pShh-1 values are the same as in Fig. 1B. D) Shh::mGL cells were transfected with dCas9-VPR, targeting the *Shh* promoter (pShh-1 guide), and another single gRNA (x axis) targeting pShh or regions upstream of Shh, and measured by flow cytometry to determine mGreenLantern MFI and normalised to pShh-1 only. E) Gating strategy used in the flow cytometry analyses, exemplified here by one replicate of the SBE2 x100 gRNA pool. Cells (top left), and singlets (top right) were gated using forward and side scatter signals to analyse only living single cells. Transfected cells were gated using the EBFP2 and mCherry signal (bottom left, see arrow). Note that in some experiments transfected cells are selected solely based on one fluorophore. Of the transfected cells, distributions of the mGreenLantern signals were used to derive the median (bottom right). F) UCSC browser map of the location of the two x50 gRNA guide pools targeted 206kb upstream of *Shh*. G) Distribution of gRNAs in plasmid pools based on perfect match read assignments from whole-plasmid sequencing data. Values in B were statistically compared to Empty/NT with t-tests and BH-corrected. Values in D were compared to 1 with one-sided t-tests and BH-corrected. Adjusted p-values <0.1 (†) and <0.05 (*) are shown (see Supplemental table 1 for statistics and replicate numbers).

**Supplemental Figure 3.**
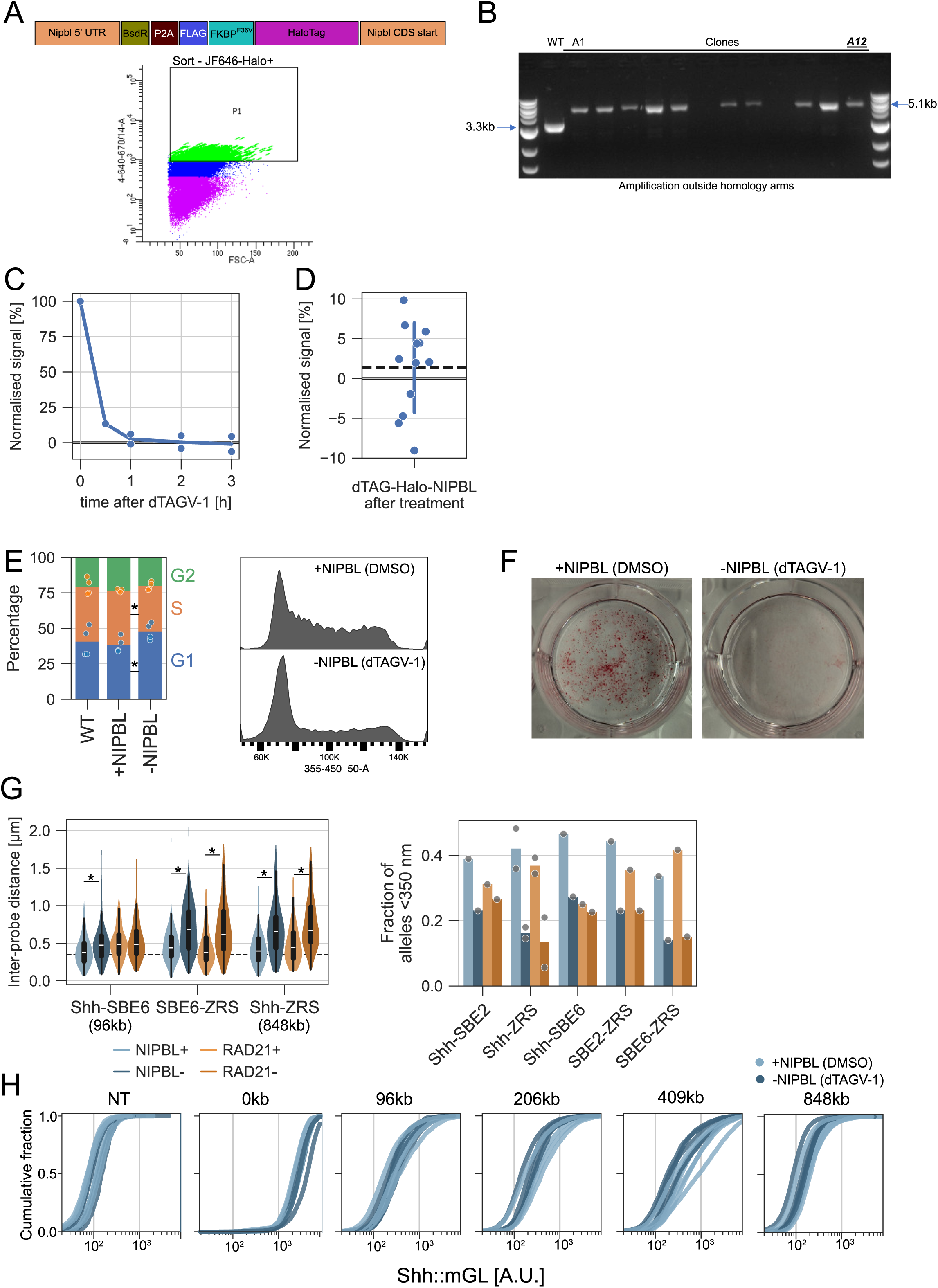
Related to Figure 2. A) Top: Schematic of the dTAG-NIPBL allele. Bottom: Sorting gate used to select HaloTag-positive cells. B) Genotyping for dTAG-NIPBL knock-in by PCR amplification from outside the homology arms surrounding the integration. The absence of a lower 3.3kb band shows homozygous targeting. The underlined clone A12 was selected for use. C) HaloTag signal based on HaloTag-JF646 flow cytometry MFI upon addition of 250 nM dTAGV-1 at different time points, min-max normalised to stained WT and untreated. D) HaloTag signal based on HaloTag-JF646 flow cytometry MFI upon addition of ≥250 nM dTAGV-1 for ≥24 hours, min-max normalised to stained WT and untreated. Dashed line indicates the mean. E) Left: Distribution of cells in G1/S/G2 cell cycle phase based on flow cytometry Hoechst signal gating for WT cells, and dTAG-NIPBL cells treated with DMSO (+NIPBL) and 250 nM dTAGV-1 (-NIPBL) for 48 hours. Values were normalised to total in those gates. Right: Example flow cytometry Hoechst signal. F) Alkaline Phosphatase staining of dTAG-NIPBL cells seeded at low density and left to clonally expand for 7 days. G) Left: Second replicate DNA-FISH data for dTAG-NIPBL cells treated with DMSO (+NIPBL) or dTAGV-1 (-NIPBL) for 24 hours or RAD21-AID cells untreated (+RAD21) or treated with Auxin for 6 hours (-RAD21). Dotted line indicates 350 nm. Right: Fraction of alleles within 350 nm for the same conditions, including replicate in Fig. 2C. H) Cumulative distribution of single-cell mGreenLantern fluorescence values for dTAG-NIPBL cells transfected with dCas9-VPR and different gRNA or gRNA pools and treated with DMSO or dTAGV-1 for 48 hours. Each line is a replicate. Values in E were statistically compared between WT and +NIPBL, and -NIPBL and +NIPBL, with paired t-tests and BH-corrected. Values in G were statistically compared between +NIPBL and -NIPBL, and +RAD21 and -RAD21 with Mann-Whitney U tests and BH-corrected. Adjusted p-values <0.1 (†) and <0.05 (*) are shown (see Supplemental table 1 for statistics and replicate numbers).

**Supplemental Figure 4.**
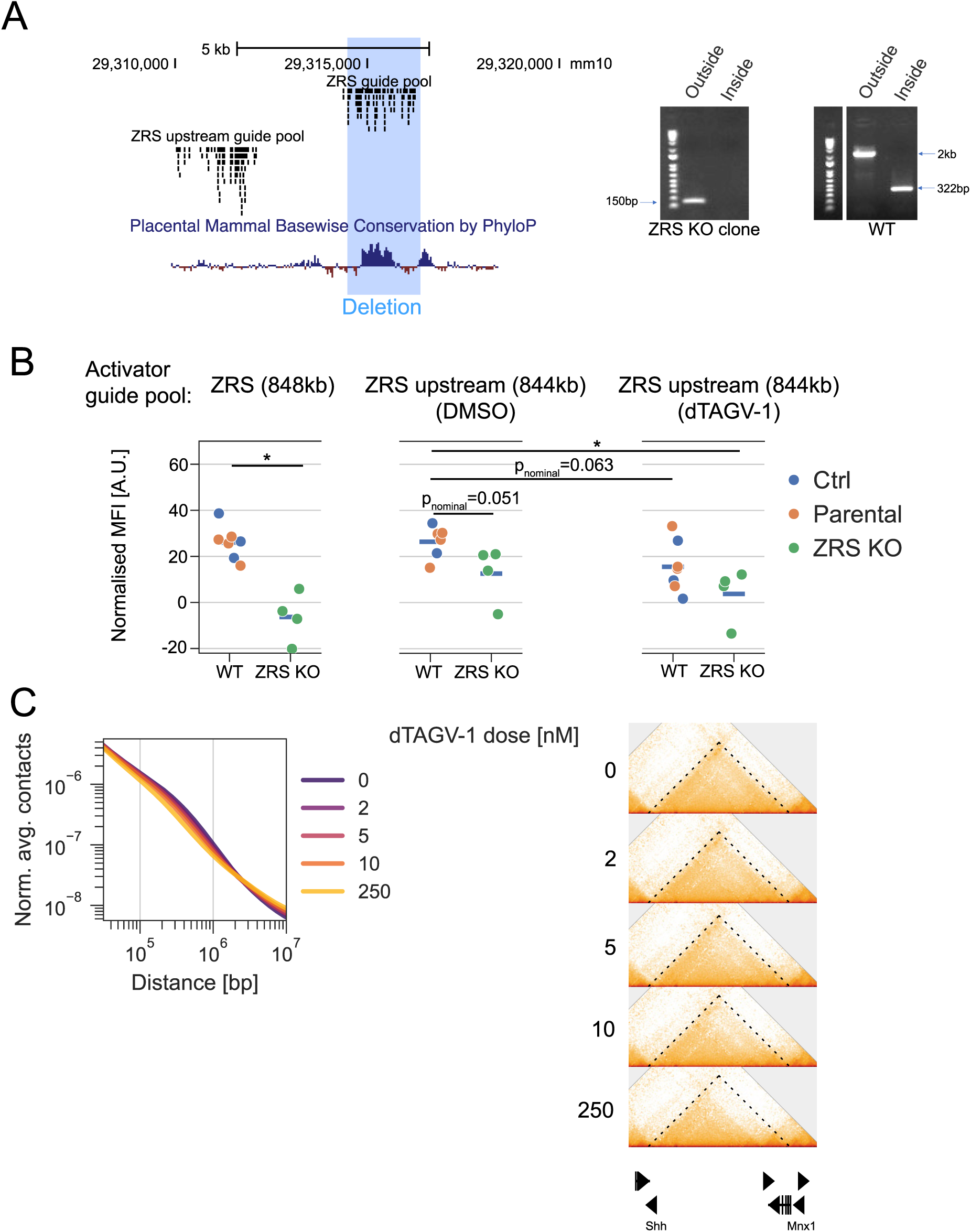
Related to Figure 2. A) Left: UCSC browser map of the location of ZRS pool and the upstream pool used in experiments in panel B. The blue highlight represents the ZRS genetic deletion. Right: Knockout verification DNA gels with amplification from outside the deleted region (left amplicon) and inside the deleted region (right amplicon). Note that the right gel was cropped to remove intervening unrelated bands. B) MFI for mGreenLantern fluorescence, normalised by subtracting the value of NT-transfected cells for cells transfected with dCas9-VPR and different gRNA or gRNA pools and treated with DMSO or dTAGV-1 for 48 hours. Left: ZRS guide pool (untreated). Middle: ZRS upstream guide pool, with DMSO treatment. Right: ZRS upstream guide pool, with dTAGV-1 treatment. WT cells represent either the parental dTAG-NIPBL clone or a control clone thereof with unedited ZRS. C) Left: P(s) curves derived from RCMC data for dTAG-NIPBL cells treated with DMSO or different doses of dTAGV-1 (nM) for 24 hours. Average contacts at the given genomic separations (bp) were normalised by the area under the curve. Right: RCMC data for the same dTAG-NIPBL cells treated with DMSO or different doses of dTAGV-1. Values in B were statistically compared between with independent t-tests and BH-corrected. Adjusted p-values <0.1 (†) and <0.05 (*) and nominal p-values <0.1 are shown (see Supplemental table 1 for statistics and replicate numbers).

**Supplemental Figure 5.**
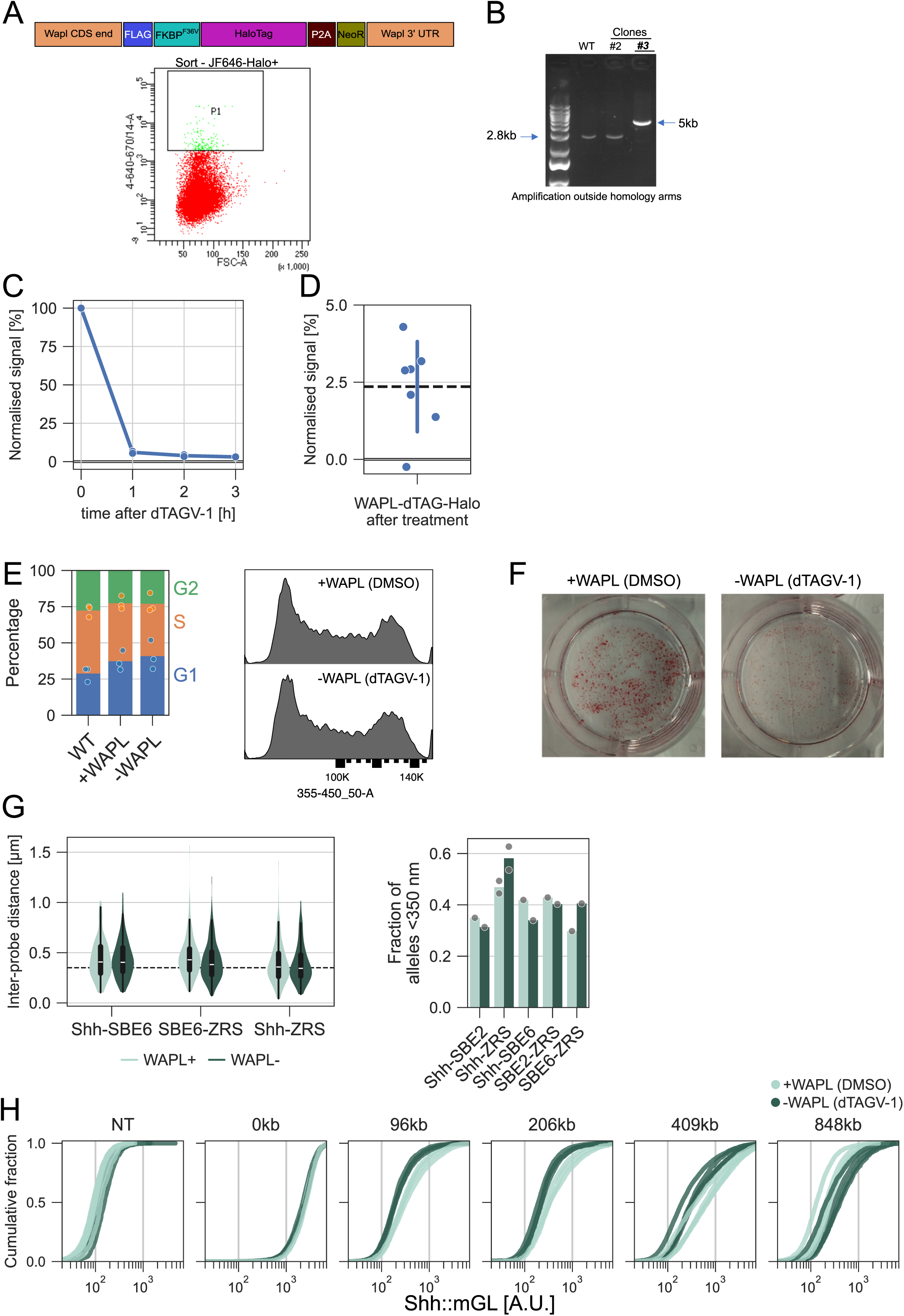
Related to Figure 3. A) Top: Schematic of the WAPL-dTAG allele. Bottom: Sorting gate used to select HaloTag-positive cells. B) WAPL-dTAG verification DNA gels with amplification from outside the homology arms surrounding the integration. The absence of a lower 2.5kb band shows homozygous targeting. The underlined clone #3 was selected for use. C) HaloTag signal based on HaloTag-JF646 flow cytometry MFI upon addition of 250 nM dTAGV-1 at different time points, min-max normalised to stained WT and untreated. D) HaloTag signal based on HaloTag-JF646 flow cytometry MFI upon addition of ≥250 nM dTAGV-1 for ≥24 hours, min-max normalised to stained WT and untreated. Dashed line indicates the mean. E) Left: Distribution of cells in G1/S/G2 cell cycle phase based on flow cytometry Hoechst signal gating and normalised to total in those gates for WT cells, and WAPL-dTAG cells treated with DMSO (+WAPL) and 250 nM dTAGV-1 (-WAPL) for 48 hours. Right: Example flow cytometry Hoechst signal. F) Alkaline Phosphatase staining of WAPL-dTAG cells seeded at low density and left to clonally expand for 7 days. G) Left: Second biological replicate of DNA-FISH data for WAPL-dTAG cells treated with DMSO (+WAPL) or dTAGV-1 (-WAPL) for 24 hours. Dotted line indicates 350 nm. Right: Fraction of alleles within 350 nm for the same conditions, including replicate in Fig. 3C. H) Cumulative distribution of single-cell fluorescence (mGreenLantern) values for WAPL-dTAG cells transfected with dCas9-VPR and different gRNA or gRNA pools and treated with DMSO or dTAGV-1 for 48 hours. Each line is a replicate. Values in E were statistically compared between WT and +WAPL, and -WAPL and +WAPL, with paired t-tests and BH-corrected. Values in G were statistically compared between +WAPL and -WAPL with Mann-Whitney U tests and BH-corrected. Adjusted p-values <0.1 (†) and <0.05 (*) are shown (see Supplemental table 1 for statistics and replicate numbers).

**Supplemental Figure 6.**
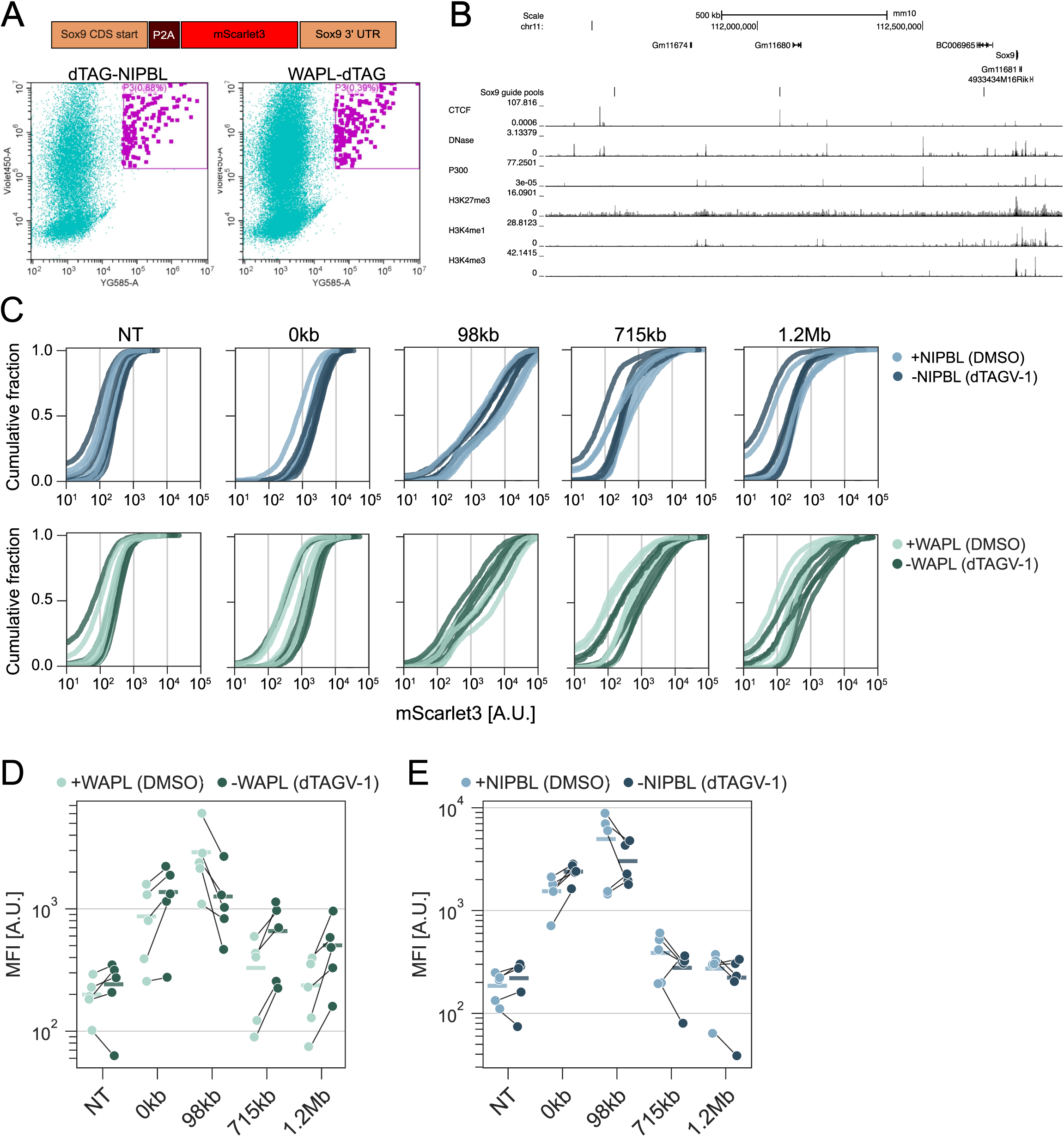
Related to Figure 4. A) Top: Schematic of the Sox9-P2A-mScarlet3 allele. Bottom: dTAG-NIPBL and WAPL-dTAG cells transfected with Cas9, gRNA and the Sox9-P2A-mScarlet3 repair template, recovered, and subsequently transfected with dCas9-VPR and guides targeting the *Sox9* TAD. Sorting gates used to select mScarlet3-positive cells are shown. B) UCSC Genome Browser tracks at the *Sox9* locus (mm10 genome), together with coordinates for the targeted gRNA pools and ENCODE mESC tracks for CTCF ChIP-seq, DNase-seq, and H3K27me3, P300, H3K4me1, and H3K4me3 ChIP-seq [141]. C) Cumulative distribution of single-cell mGreenLantern fluorescence values for dTAG-NIPBL and WAPL-dTAG *Sox9* reporter cells transfected with dCas9-VPR and different gRNA or gRNA pools and treated with DMSO or dTAGV-1 for 48 hours. Each line is a replicate. D-E) As in panel C, but showing median fluorescence intensities per replicate.

**Supplemental Figure 7.**
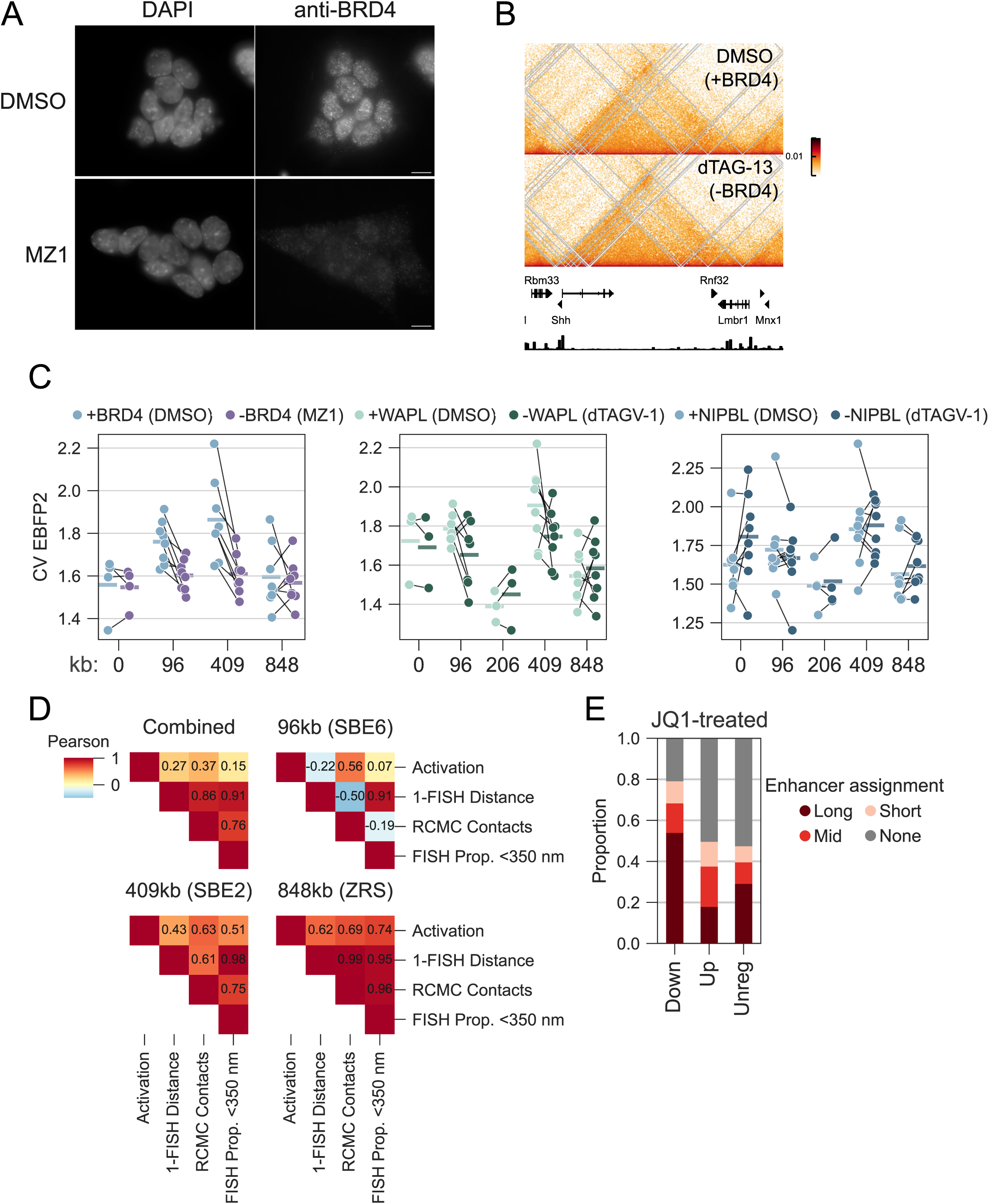
Related to Figures 5 and 6. A) Immunofluoresence images of mESCs grown in the presence of DMSO or 100 nM MZ1, stained against DNA (DAPI) and BRD4 (anti-BRD4). Scale bar = 5 microns. B) Hi-C data for dTAG-BRD4 cells from Linares-Saldana et al. [92] treated with DMSO (top) or dTAG-13 (bottom) for 24 hours. C) Coefficient of variation (CV) of the EBFP2 signal (transcribed from the transfected plasmid) of the transfected population upon depletion of BRD4, WAPL, and NIPBL. D) Pairwise Pearson correlation coefficients for *Shh* activation (normalised MFI), inverse median FISH distances (1 minus median; μm), RCMC contacts between *Shh* and the activation site (log contacts), and the proportion of FISH alleles within 350 nm. Combined includes the three activation sites with all data types (96kb, 409kb, and 848kb). E) Distribution of gene categories in the Down-/Up-/Unregulated sets for cells treated with JQ1, as in Fig. 6 (data from Nakato et al. [43]).

**Supplemental Figure 8.**
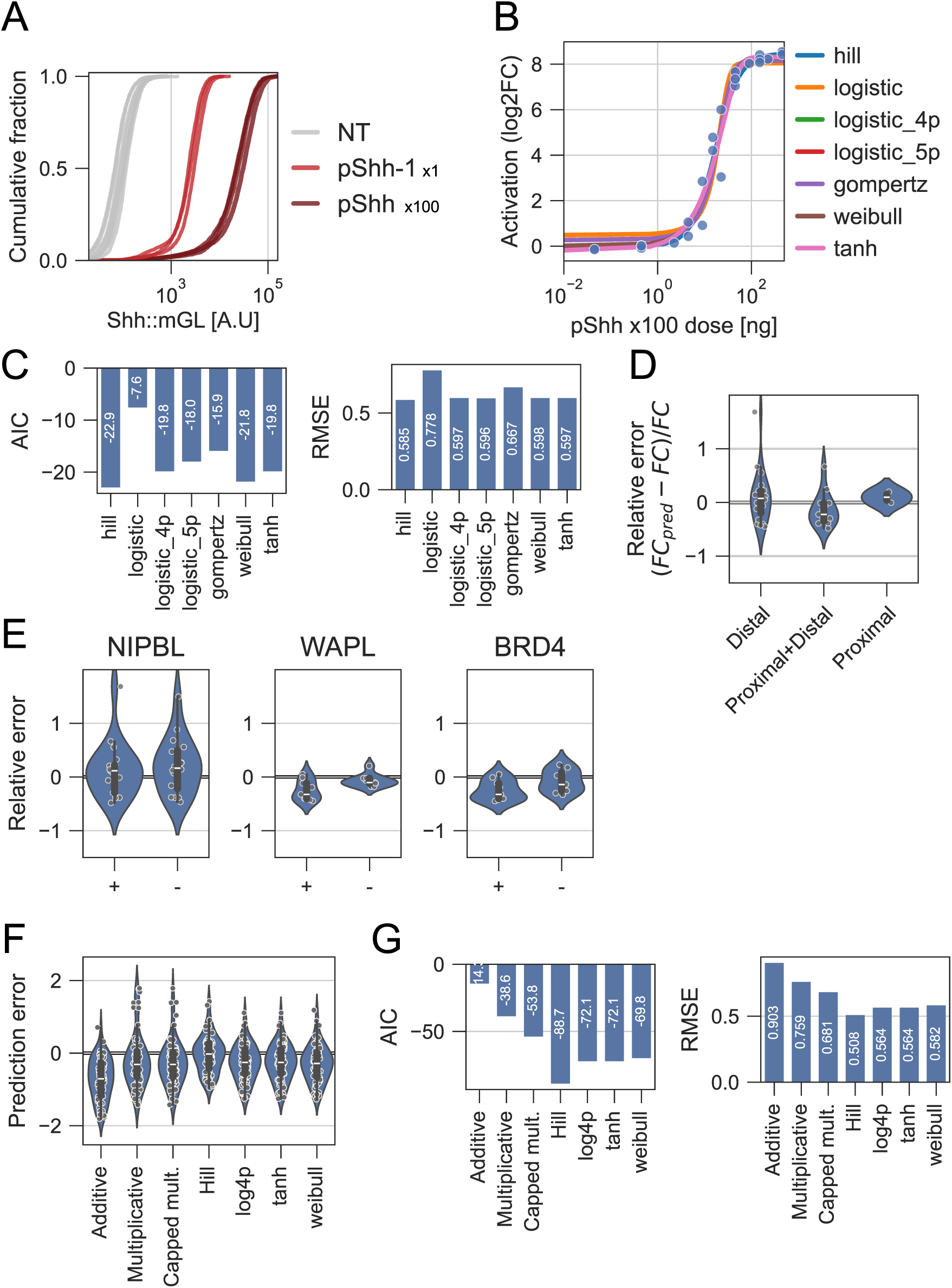
Related to Figure 7. A) Shh::mGL reporter cells were transfected with dCas9-VPR a non-targeting gRNA (NT), a promoter-targeting gRNA (pShh-1) or a gRNA pool targeting the promoter (pShh x100). Transfected cells were analysed for mGreenLantern fluorescence. Cumulative distribution of single-cell fluorescence values are shown. Each line is a replicate. B) MFI for mGreenLantern fluorescence normalised to NT-transfected cells by division (points, log2 fold-change; y axis) for Shh::mGL reporter cells transfected with dCas9-VPR and different doses [ng] of a gRNA pool targeting the Shh promoter (x axis). Various sigmoidal functions were fitted to the data. C) Akaike information criteria (AIC) and root mean square error (RMSE) metrics for the fitted functions in panel B. D) Violins show the relative error (error normalised by magnitude) of the response model prediction of dual activation (see Fig. 7) using pairs of gRNAs targeting both distally, one proximal and one distal, or both proximally. Boxes show interquartile range (Q1-Q3) and whiskers show remaining data range. E) Violins show the relative error of the response model prediction of dual activation upon depletion of NIPBL, WAPL, and BRD4, where + and – represent presence or absence of the protein (i.e. + shows control). Boxes show interquartile range (Q1-Q3) and whiskers show remaining data range. F) Violins show the prediction errors for the various models used to predict dual activation using all data. Boxes show interquartile range (Q1-Q3) and whiskers show remaining data range. G) Akaike information criteria (AIC) and root mean square error (RMSE) metrics for the prediction of dual activation in panel F.

## Table legends

**Supplemental table 1. p-values**

Description of the comparisons (Condition), statistical test (test), replicate numbers (N), and p-values (pval). Adjusted p-values (padj) are after FDR-correction of all values in one Source, and sig indicates padj<0.05.

**Supplemental table 2. Plasmids**

Description of plasmids used in the study and their sources.

**Supplemental table 3. Primers/oligonucleotides**

All oligos used for cloning, cell line verification, RCMC library preparation, and individual gRNA cloning.

**Supplemental table 4. gRNA pools**

Ordered gRNA oPools, including overhangs for amplification.

**Supplemental table 5. BACs**

BACs used in the RCMC protocol. Coordinates in mm10.

**Supplemental table 6. External data**

All data from external sources used in the study.

## Data availability

RCMC data have been uploaded to GEO (GSE329685). Flow cytometry summary statistics, DNA-FISH data, and custom analysis scripts are available at https://github.com/efriman/Friman_etal_2026_LoopCoop. Published data used are designated in Suppl. Table 6. All data and code are available upon request.

## Acknowledgements

This work was supported by UKRI Medical Research Council (MRC) University Unit grant MC_UU_00035/7 and European Research Council (ERC) synergy grant GeneMotors. H.J., G.A., A.M.G and G.W. were supported by an MRC Doctoral Training Programme. This work has made use of the resources provided by the Edinburgh Compute and Data Facility (ECDF; http://www.ecdf.ed.ac.uk). We want to thank Lizzie Freyer and Michael Rennie at the IGC Flow Cytometry Service for their help.

## Author contributions

E.T.F. and W.A.B. conceptualised the project, supervised the work, and wrote the first manuscript draft. E.T.F. performed data analyses, visualizations, and immunofluorescence experiments. E.T.F., H.J.J., G.A., and G.W. performed flow cytometry experiments. E.T.F., H.J.J., G.A., and A. M. generated cell lines. H.J.J. and L.G.A. performed RCMC experiments. S.B. performed DNA-FISH, microscopy, and image analysis. E.T.F. H.J.J., G.A., M.F., and G.W. generated plasmid constructs. M.F., A.M.G., I.W., and L.L performed important preliminary experiments not used in the final manuscript. W.A.B. acquired funding. All authors provided feedback on the manuscript and approved its contents.

## Notes

### Competing Interest Statement

The authors have declared no competing interest.

