## Supplemental protocol for "Loop extrusion dynamics and cooperative activation independently regulate enhancer-driven transcription"

**Cloning gRNA pool guides into U6-sgRNA BbsI expression plasmids**

1. Designing guides

Go to <http://crispor.tefor.net/> or <http://crispor.gi.ucsc.edu/> and enter a DNA sequence surrounding your region of interest (e.g. enhancer/promoter), including extra flanks (up to ~2kb to get at least 100 good guides). Select your genome on step 2 and use default settings on step 3 (20bp-NGG-SpCas9).

Go to the bottom of the table and download “Guides” and “Off-target” in the links next to “tab-sep format”. Use the python script example (<https://github.com/efriman/Friman_etal_2026_LoopCoop/blob/main/GuidePools/pShhx100_gRNApool_design.ipynb>) to filter out and select the top 100 guides cloned with overhangs as below:

ATGAGTCCATCAGGAAGACGGCACCNNNNNNNNNNNNNNNNNNNNGTTTGGGTCTTCGCTGAGCATGATT

Order your ≥100 guides as an IDT oPool at the smallest scale (1 pmol/oligo)

2. Digest plasmid (can be scaled down)

- 5 ug pU6-BbsI plasmid (e.g. pU6sgEnhBFP, pSPgRNA, pX330-based)
- 1 ul BbsI-HF
- 5 ul CutSmart
- H2O to 50 ul

Incubate 37C for >1 hour. Run on ~0.8% agarose and cut out single band and gel purify using Qiagen Gel Extraction Kit or equivalent. Measure concentration on Nanodrop. Can be stored in -20C.

3. Amplify guides (can be scaled down)

Resuspend the ordered oPool in 150 ul buffer EB or equivalent (avoid excess EDTA) and amplify:

- 2 ul resuspended oPool
- 100 ul KAPA HiFi HotStart ReadyMix 2x
- 200 nM gRNA pool F (ATGAGTCCATCAGGAAGACGGCACC)
- 200 nM gRNA pool R (AATCATGCTCAGCGAAGACCCAAAC)
- H2O to 200 ul

Split into 2xPCR tubes (100 ul each) and amplify using 8 cycles: 95C 3m-->(98C 20s-->65C 15s-->72C 10s)x8-->72C 1m. Run 25 ul on an agarose gel (>1%) to confirm one product at 70 bp. Purify the rest using a QIAgen MinElute Reaction Cleanup Kit column or equivalent (N.B. some kits will purify away <100bp fragments). Measure concentration on Nanodrop (should be ~20-50 ng/ul in 10 ul elution). Can be stored in -20C.

4. Digest PCR product

- All product from step 2 (~8 ul after elution and Nanodrop)
- 2 ul CutSmart
- 0.5 ul BbsI-HF
- 9.5 ul H2O (20 ul total)

Incubate at 37C for >1 hour. Can be stored in -20C.

5. Ligation

- 1 ul digested PCR product (from step 4)
- ~20 ng pU6 plasmid digested with BbsI (from step 2)
- 0.5 ul T4 DNA ligase buffer
- 0.5 ul T4 DNA ligase
- 0.25 ul BbsI-HF (optional but recommended)
- H2O to 5 ul total

Incubate at least 1 hour at RT.

6. Transformation and culture

- Thaw library efficiency DH5a competent cells (cat. 18263012) or other high efficiency bacteria on ice
- Mix 50-100 ul cells with 5 ul ligation reaction (step 5) in ice-cold Eppendorf or PCR tube. Pipet up and down as little as possible
- Leave 30 min on ice
- Heat shock 42C 30 seconds and place back on ice for ~2 min
- Add 500 ul SOC and incubate with rotation 37C 1 hour
- Plate 10 ul of transformation onto pre-warmed LB+Amp plates (dilute with 200 ul SOC to help spread out), incubate 37C overnight
- Take the rest of the transformation volume and transfer to 30 ml LB+Amp (at least room temp) liquid culture. Incubate at 37C with rotation overnight
- Next day, count the number of colonies on the plate and estimate coverage. For example, if there are 100 colonies from 50 ul original starting cells diluted in 500 ul SOC: 50 colonies/10ul*600ul=3000 ~ 30X coverage for 100-guide library.
- Spin down liquid culture >5000xg 15 min and midiprep (or save pellet in -20C for later)
- Optional: Inoculate 3-5 colonies from plate into 2 ml LB+Amp cultures and incubate overnight 37C with rotation. Next day, miniprep and Sanger sequence with a U6 forward primer (e.g. ACTATCATATGCTTACCGTAAC) to confirm correct inserts (guides should be where N: acgaaacaccNNNNNNNNNNNNNNNNNNNNgttttagagct)
  - If a large proportion (>1/5) of your colonies are undigested plasmid without guide, redo the digestion (step 2) longer and/or separate longer on gel
  - If you have mismatches in your cloned guides, considering lowering the number of cycles in step 3
- Sequence the pool using Plasmidsaurus or equivalent whole plasmid sequencing service and compare inserts to your expected oPool using the script (<https://github.com/efriman/Friman_etal_2026_LoopCoop/blob/main/GuidePools/GuidePool_assignment.ipynb>)
